# Screening the MMV Pathogen Box reveals the mitochondrial *bc*_1_-complex as a drug target in mature *Toxoplasma gondii* bradyzoites

**DOI:** 10.1101/2023.11.23.568420

**Authors:** Deborah Maus, Elyzana Putrianti, Tobias Hoffmann, Michael Laue, Frank Seeber, Martin Blume

**Affiliations:** P 6: Metabolism of Microbial Pathogens, Robert Koch Institute, 13353, Berlin, Germany; FG 16: Mycotic and Parasitic Agents and Mycobacteria, Robert Koch Institute, 13353, Berlin, Germany; ZBS 4: Advanced Light and Electron Microscopy, Centre for Biological Threats and Special Pathogens 4, Robert Koch-Institute, 13353, Berlin, Germany

**Keywords:** Apicomplexa, in vitro tissue cysts, *bc*_1_-complex, metabolomics, mitochondrial metabolism

## Abstract

The apicomplexan parasite *Toxoplasma gondii* infects 25-30% of the global human population and can cause life-threatening diseases in immunocompromised patients. The chronically infectious forms of the parasite, bradyzoites, persist within cysts in brain and muscle tissue and are responsible for its transmission and remission of the disease. Currently available treatment options are very limited and are only effective against the fast-replicating tachyzoites but fail to eradicate the chronic stages of *T. gondii*. The cause of these treatment failures remains unclear. Here, we utilized our recently developed human myotube-based culture model to screen compounds from the MMV Pathogen Box against pan-resistant in vitro bradyzoites and identified multiple compounds with simultaneous activity against tachyzoites and bradyzoites. Stable isotope-resolved metabolic profiling on tachyzoites and bradyzoites identified the mitochondrial *bc*_1_-complex as a target of bradyzocidal compounds and defines their metabolic impacts on both parasite forms. Our data suggest that mature bradyzoites rely on mitochondrial ATP production.

## Introduction

*Toxoplasma gondii* is an ubiquitous apicomplexan parasite that infects virtually all warm-blooded animals, including 25-30 % of humans (Montazeri et al. 2020). Of all food-borne pathogens, *T. gondii* causes 20% of the total disease burden in Europe (Havelaar et al. 2015). Although most acute infections caused by the fast replicating tachyzoite stage of the parasite remain asymptomatic in immunocompetent individuals, the parasite persists life-long in the form of encysted semi-dormant bradyzoites (Weiss and Dubey 2009). Currently, no medical treatment exists to eradicate these tissue cysts with their rigid cyst wall from infected individuals. *T. gondii* is transmitted through undercooked meat products of infected livestock but also via feline-shed oocysts that can contaminate water sources and soil (Weiss and Dubey 2009). Severe toxoplasmosis occurs after reactivation of cysts or primary infection of immunosuppressed individuals or fetuses of seronegative mothers and can lead to systemic infections, uveitis, encephalitis as well as congenital transmission with potentially fatal outcomes (Montoya and Liesenfeld 2004). Hence, tissue cysts of *T. gondii* constitute a long-term source for transmission and risk of acute disease. Also, other related cyst-forming coccidian parasites such as *Neospora caninum* and *Besnoitia besnoiti* pose an immense veterinary burden by causing abortion mainly in sheep and severe chronic infections in cattle and horses, respectively. Current treatment options exist only against the tachyzoite form of these parasites and include antifolates, such as pyrimethamine or trimethoprim, that inhibit the bifunctional dihydrofolate reductase-thymidylate synthetase, in combination with sulfadiazine, which blocks the dihydropteroate synthetase. These enzymes are implicated in DNA synthesis in both the parasite and side-effects often require pre-mature termination of the treatment (Katlama, De Wit, et al. 1996; Alday and Doggett 2017). Another potent drug target in coccidians and other apicomplexan parasites such as *Plasmodium* spp. is the cytochrome *bc*_1_-complex in the mitochondrial respiratory chain, which is inhibited by a number of approved drugs, such as the naphthoquinones atovaquone (ATQ) (Meneceur et al. 2008; Nixon et al. 2013) and buparvaquone (BPQ) (Sharifiyazdi et al. 2012). However, both coenzyme Q analogs including bioavailable nanosuspensions (Shubar et al. 2011) fail to eradicate encysted bradyzoites of *T. gondii* and *N. caninum* from the brains of infected mice after multiple weeks of treatment (Doggett et al. 2012; Müller et al. 2015; Doggett et al. 2020). Experimental inhibitors of the *bc*_1_-complex includes 1-hydroxy-2-dodecyl-4(1H)quinolone (HDQ) that arrests the growth of asexual blood stages of the related malaria parasite *P. falciparum* (Vallières et al. 2012). However, in *T. gondii* HDQ appears to inhibit multiple mitochondrial election donors such as the alternative NAD-dehydrogenase and the dihydroorotate dehydrogenase in tachyzoites (S. S. Lin, Gross, and Bohne 2009). It does not affect mature bradyzoites in vitro (Christiansen et al. 2022), and HDQ derivatives do not clear *T. gondii* cysts from the brain of infected mice (Bajohr et al. 2010). In addition, endochin-like quinolones constitute a promising series of potent *bc*_1_-complex inhibitors that are active against *T. gondii* (McConnell et al. 2018) and *Plasmodium* parasites (Song et al. 2018). However, these compounds also do not fully eradicate the cysts *in vivo* (Doggett et al. 2012, 2020). Similarly, bumped kinase inhibitors that selectively inhibit parasite calcium-dependent protein kinases in various apicomplexans fail to clear tissue cysts *in vitro* (Christiansen et al. 2022) and *in vivo* (Hulverson et al. 2019). Despite these efforts, it remains unclear whether the target proteins of these drugs are rendered dispensable for the survival of bradyzoites or whether the eradication failures have other reasons, such as poor permeability across the blood brain barrier and/or the cyst wall. To date, the genetic tools available to directly characterize the function of essential genes in bradyzoites are limited to promoter replacements (Smith et al. 2021), while no genetic methods exist to directly interrogate the fitness contribution of mitochondrially encoded genes such as the cytochrome b subunit of the cytochrome *bc*_1_-complex (J.-L. Wang et al. 2016). Consequently, reverse pharmacological approaches, that involve screening of inhibitors against target proteins, remain difficult and the underlying reasons for limited *in vivo* efficacies remain uncertain.

Here we aim to address these shortcomings and identify essential processes in bradyzoite by a combination of a phenotypic screen and subsequent metabolomics-driven identification of the modes of actions of hit compounds. We employed a recently developed human myotube-based culture system to generate mature *T. gondii* drug-tolerant bradyzoites (Christiansen et al. 2022; Maus et al. 2024) to screen the MMV Pathogen Box. These *in vitro* cysts exhibit resistance against antifolates, bumped kinase inhibitors and HDQ. They also express hallmarks of *in vivo* cysts including oral infectivity and a cyst wall (Christiansen et al. 2022). The Medicines for Malaria Venture (MMV) Pathogen Box provides a curated set of bioactive molecules with activity against a number of pathogens including *P. falciparum*, *Cryptosporidium parvum*, *Schistosoma spp*. and diseases caused by kinetoplastids (Veale 2019). Untargeted and stable isotope-resolved metabolomic analyses of active drugs identified established and unsuspected modes of action in tachyzoites and bradyzoites, respectively. Our data directly show metabolic consequences of *bc_1_*-complex inhibition in bradyzoites and suggest its crucial role in ATP production.

## Material and Methods

### Tachyzoite Culture

Prugeniaud Δhxgprt tdTomato (John et al. 2009) (PruTom) and RH-Rep1/2-S9(33-159)-GFP (DeRocher et al. 2000; Thomsen-Zieger, Schachtner, and Seeber 2003) (RH-S9) parasites were maintained in human foreskin fibroblasts (HFF) cells (BJ-5ta human foreskin fibroblasts ATCC, CRL-4001) with Dulbecco’s modified Eagle’s medium (DMEM) containing 25 mM glucose, 4 mM L-glutamine, 1 mM sodium pyruvate, 100 U/mL penicillin, 100 µg/mL streptomycin and 1 % heat-inactivated fetal bovine serum (FBS, Gibco) at 37°C with 10 % CO_2_ (Christiansen et al. 2022).

### *In vitro T. gondii* tissue cyst culture

The *T. gondii in vitro* cyst culture was performed as in (Christiansen et al. 2022; Maus et al. 2024). In short, KD3 myoblasts (Shiomi et al. 2011) were grown to 70 % confluency, followed by the induction of myotube differentiation by changing the medium to DMEM as described above but with 2% horse serum instead of FBS and additionally supplemented with 10 µg/mL human insulin, 5 µg/mL human holo-transferrin and 1.7 ng/mL Na_2_SeO_3_. After five days, the myotubes were infected with PruTom (MOI 0.1) in bicarbonate-free Roswell Park Memorial Institute medium (RPMI) containing 5.5 mM glucose, 50 mM HEPES, 4 mM L-glutamine, 100 U/mL penicillin, 100 µg/mL streptomycin, 2 % horse serum, 10 µg/mL human Insulin, 5 µg/mL human holo-transferrin and 1.7 ng/mL Na_2_SeO_3_ at pH 7.2. Cultures were incubated at 37 °C at ambient CO_2_ for four weeks and the medium was changed every two to three days. A comprehensive scheme of the procedure is shown in Fig. S1.

### MMV Pathogen Box Screening Procedure

All Pathogen Box compounds were stored at -20°C (Veale 2019). DMSO-solved compounds were diluted in DMEM or RPMI media and stored at 4°C until use. Host cells were grown in transparent, flat-bottom 96-well plates and infected as described above to cultivate either tachyzoites or in vitro tissue cysts. One column per plate was left uninfected to estimate fluorescence background. Four hours after infection with tachyzoites or after four weeks of bradyzoite maturation the culture medium was replaced by medium containing either 0.1 % DMSO (negative control) or 10 µM of the respective Pathogen Box compound. During the seven-day treatment period the medium was renewed every two to three days. It was then changed to tachyzoite growth medium to induce re-differentiation of viable bradyzoites to tachyzoites, and then the culture was monitored over 28 days. Prior to each medium exchange, the cultures were monitored for compound crystallization. The fluorescence was measured in a fluorescence plate reader (Tecan Infinite M200 PRO) (excitation Λ = 554 nm, emission Λ = 589 nm). Fluorescence values passing a threshold of 2,000 after background subtraction indicated proliferating parasites and hence an ineffective treatment (Fig. 1).

**Figure 1:**
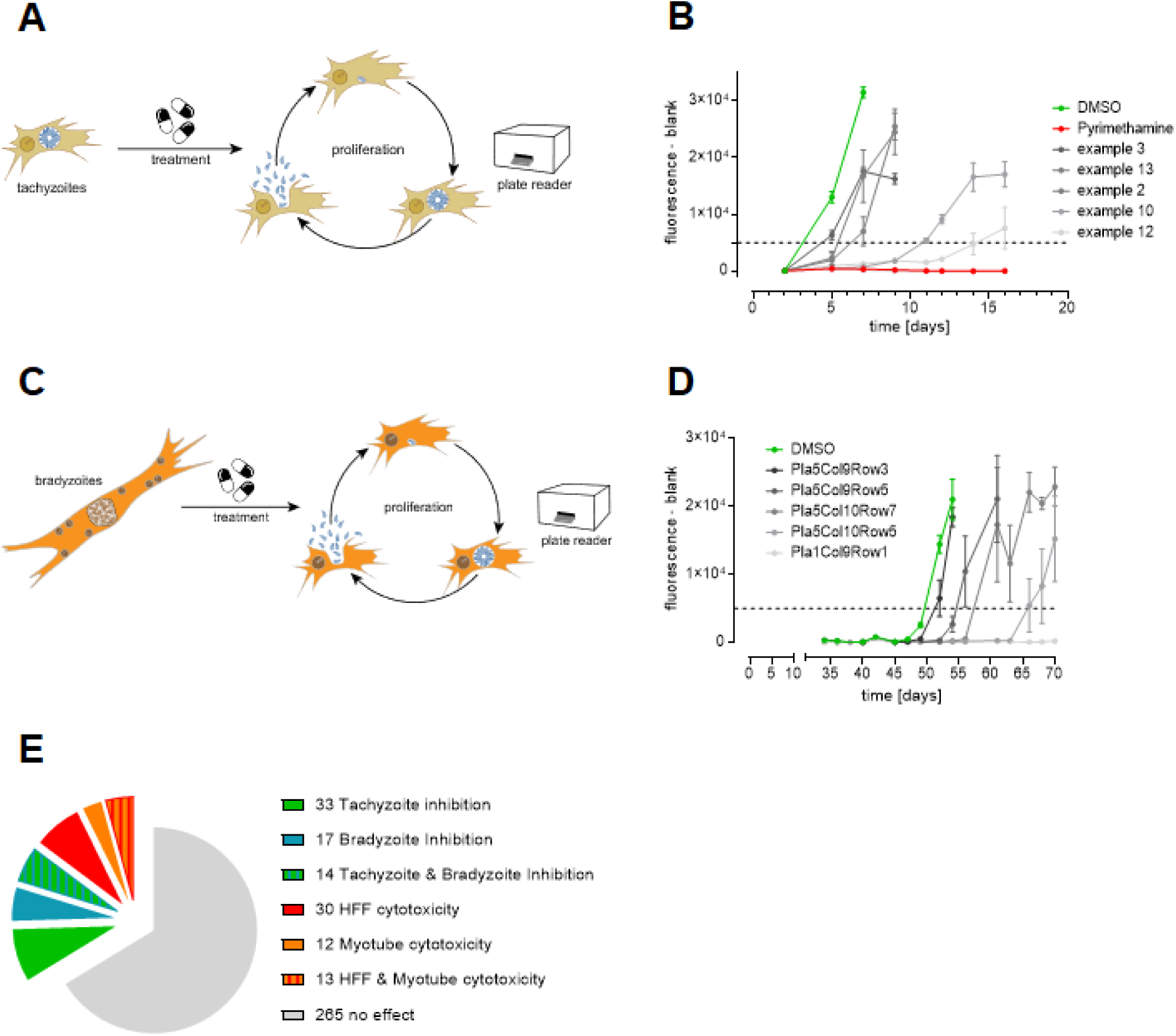
An in vitro screen reveals tachy- and bradyzocidal properties of the MMV Pathogen Box compounds. (A) Experimental scheme of the MMV compound screen on *Toxoplasma gondii* tachyzoites in human foreskin fibroblast cells (B) Growth curves of tachyzoites treated with 10µM of each MMV compound for 7 days (shades of grey), a solvent control (green), and an inhibition control (red) (C) Experimental scheme of the MMV compound screen on bradyzoites matured in myotubes (D) Growth curves of bradyzoites treated with 10µM of each compound (shades of grey) or a solvent control (green) for 7 days, followed by a tachyzoite regrowth phase (E) Pie chart displaying the number of Pathogen Box compounds and their effects on our in vitro cultures.

### Resazurin viability assay and cytotoxicity

After the growth assays were completed, the host cell monolayers of both HFF and KD3 myotubes were washed with PBS and incubated with 500 nM resazurin in Milli-Q water at 37 °C for 4 hours. Fluorescence was measured (excitation Λ = 540 nm, emission Λ = 580 nm) in a fluorescence plate reader and normalized to the average fluorescence values of the DMSO-treated controls, which were set at 100 %. During bradyzoite assays KD3 host cells were visually inspected after drug exposure for one week. After completion of the re-growth phase for four weeks the colorimetric resazurin was performed. These values were reported as lowest cytotoxicity values (Supplemental Fig 2D). During secondary tests of putatively dually active compounds resazurin assays were performed in a concentration dependent manner (Supplemental File S1, Supplemental Fig 2D).

### The dose response of dually active compounds

For the measurement of LC50 and minimal lethal concentrations tachyzoite and bradyzoite cultures were treated with the MMV compounds which were pipetted in a serial dilution across the 96-well plate and treated for 7 days. Afterwards, the compound containing medium was removed and possible regrowth was monitored for another 7 days (tachyzoite assay) or 21 to 28 days (bradyzoite assay) in a fluorescence plate reader (Tecan Infinite M200 PRO) (excitation Λ = 554 nm, emission Λ = 589 nm). The relative fluorescence units were background-subtracted. Depending on the development of the DMSO control, a late timepoint when DMSO-treated cultures still continued exponential growth was chosen to calculate the half-maximal inhibitory concentration (IC_50_) for tachyzoites. This time point was between day 4 and 7. Bradyzoite cultures were of lower MOI, and we calculated LC_50_ values between experiments from 7 to 28 days post treatment when exponential growth occurred in mock treated cultures (Fig. 2). The concentration data was log-transformed and a non-linear regression (curve fit with variable slope) was performed to determine the dose response (Prism 9, GraphPad).

**Figure 2:**
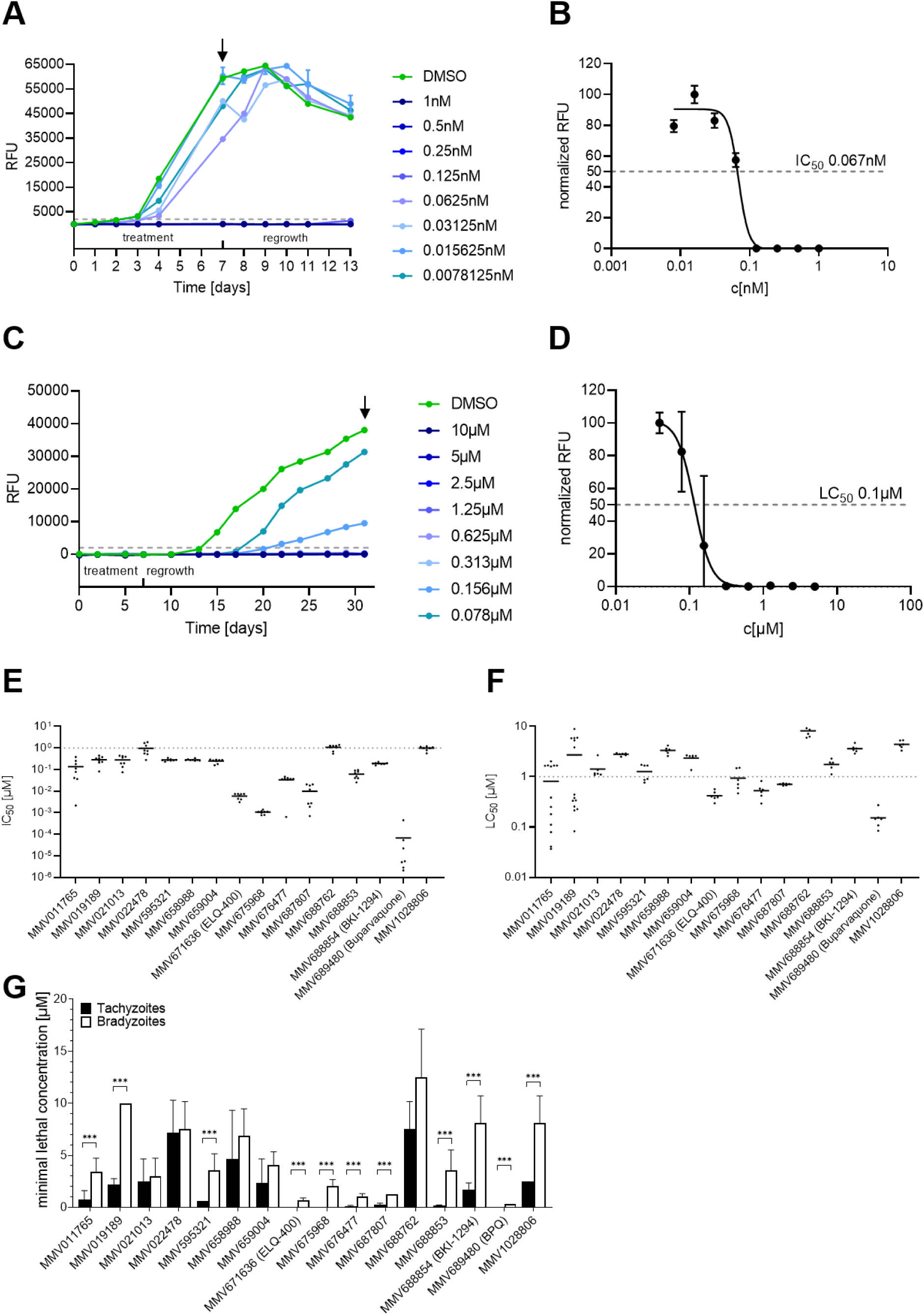
Confirmed Pathogen Box compounds that target both tachyzoites and mature bradyzoites. (A) shows the growth curve of MMV689480 (Buparvaquone), one of the 16 dually active compounds, in a tachyzoite growth assay (n=8). The infected cultures were treated for 7 days and regrowth in absence of the inhibitor was observed for another week. Depending on the development of the DMSO control, the half-maximal inhibitory concentration (IC_50_) was determined at either day 4 or 7 (arrow). (B) shown is the respective data plotted as relative fluorescence units depending on the concentration at day 7. (C) With 4 weeks old, mature bradyzoite cultures (n=6), the half-maximal lethal concentration (LC_50_) was determined at the time point which ideally reflected the dose-response curve (in the case of Buparvaquone at day 31; arrow). (D) The LC_50_ of Buparvaquone on encysted bradyzoites determined at day 31. All IC_50_ and LC_50_ of dually active compounds are summarized in (E) for tachyzoite and (F) for bradyzoite cultures. The grey dotted line indicates 1 µM in both charts. (G) Comparison of the minimal lethal concentration of each compound on tachyzoites and bradyzoites. Mann-Whitney test of eight replicative measurements on tachyzoites and six replicates of bradyzoites from two independent experiments each *** p < 0.001.

### *In silico* predictions

MMV provided the SMILES IDs for every compound, which were compiled with LightBBB (Shaker et al. 2020) to predict the blood-brain permeability. SwissADME (Daina, Michielin, and Zoete 2017) was used to predict the gastrointestinal absorption capability. The logP values (Fig. S2) were provided by MMV and remaining logP values were calculated on SwissADME (Daina, Michielin, and Zoete 2017) as consensus values.

### Fluorescence stain of mitochondrial activity

This assay was performed on tachyzoites and bradyzoites. For the tachyzoite experiment, HFF cells were infected with RH-S9 parasites at a MOI 0.1 and treated for 24 h with 1 µM atovaquone, 25 nM buparvaquone or 5 µM MMV1028806, respectively, corresponding to five-fold of the respective IC_50_ values, or 0.1% DMSO as a solvent control. For the bradyzoites experiment, 4 weeks-old ME49 *in vitro* cysts were treated for 24h with 5 µM atovaquone, 500 nM buparvaquone or 5 µM MMV1028806, respectively, corresponding to the lowest lethal concentration, or 0.1 % DMSO as a solvent control.

The assay was repeated with additional control treatments (see Supplemental Fig S5). In this experiment, HFF cells were infected with RH-S9 tachyzoites and, 4h post-infection, incubated for 24h with 0.1% DMSO (solvent control), 1 µM atovaquone, 25 nM buparvaquone, 5µM MMV1028806 (MMV), 1 µM pyrimethamine (Pyri), 10 nM clindamycin, or 25 µM 6-diazo-5-oxonorleucine (DON). Concentrations reflected the five-fold IC_50_ values, except for DON, which does not inhibit growth.

Subsequent staining and imaging procedures were identical across all experiments. Cultures were then stained with 500 nM Mitotracker Deep Red FM (MT; Thermo Fisher) in DMEM for 40 min at 37 °C, followed by a 20 min de-staining with DMEM and final fixation with 1 % paraformaldehyde for 20 min at room temperature. The cyst wall was decorated with *Dolichos biflorus* Agglutinin (DBA) and stained with Streptavidin-Cy2. Mowiol containing DAPI (1:3,000) was used to mount the cells. To avoid a biased image acquisition, all parasite vacuoles or cysts identified in a specimen along a horizontal axis were imaged. Tachyzoite images were taken with a Zeiss Axio Imager Z1 Microscope equipped with the ApoTome 2 at identical exposure times (DAPI 200 ms, MT 500 ms, S9-GFP 200 ms, Brightfield 10 ms) using a Zeiss Plan APOCHROMAT 63x/1.4 oil objective lens. Bradyzoite images were acquired on a Leica Stellaris 8 laser scanning confocal microscope using a Leica HC PL APO CS2 63x/1.4 oil objective lens. The pinhole diameter was set to 1 AU. All detectors were operating in photon counting mode. The acquisition parameters were set as follows. DAPI: diode laser at 405nm_exc_ with 2.2%, 430-550nm_em_ with a HyD S-detector. Cy2: WLL at 468nm_exc_ with 8.5%, 473-591nm_em_ with a HyD S-detector. Mitotracker: WLL at 641nm_exc_ with 2.7%, 650-784nm_em_ with a HyD X-detector (4x line average). The zoom was set to 7x, images were acquired in frame-sequential mode with a unidirectional scan speed of 400 Hz, the pixel dwell time for all channels was 3.18µs. For z-Stacks, z-step size was set to 0.299 µm.

Using ImageJ (Schindelin et al. 2012), the intensity threshold of individual pixels of MT (190 to 255 for tachyzoites and 125 to 65535 for bradyzoites) and DAPI (140 to 255 for tachyzoites and 363 to 65535) channels was cut off and the signal intensity was calculated per parasitophorous vacuole or cyst.

### Oxygen consumption assay

To assess mitochondrial oxygen consumption in *T. gondii* RHΔku80 tachyzoites, we employed the MitoXpress Xtra Oxygen Consumption Assay (Agilent Technologies, Cat# MX-200-4) following the manufacturer’s high-sensitivity (HS) method with modifications. Parasites were freshly harvested upon egress and resuspended at a density of 1 × 10⁷ tachyzoites per reaction in phenol red-free D1 medium lacking glucose but supplemented with glutamine to promote mitochondrial electron transport chain (mETC) activity via the GABA shunt (MacRae, Sheiner et al. 2012).

Parasites were incubated in medium with one of the following treatments immediately before measurement: 0.1% DMSO (vehicle control), 1 µM antimycin A (complex III inhibitor, positive control), 10 µM FCCP (uncoupler, commonly used to determine the maximum respiration capacity), 1 µM ATQ, 1 µM BPQ, or 1 µM MMV1028806. Untreated control wells contained 1 × 10⁷ tachyzoites in 0.1% DMSO. Background fluorescence was determined from wells containing MitoXpress Xtra reagent but no parasites.

Each 100 µL reaction was supplemented with 10 µL of reconstituted MitoXpress Xtra reagent. To prevent oxygen diffusion from ambient air, 100 µL of prewarmed HS mineral oil was carefully layered over each well. Plates were immediately transferred to a prewarmed Tecan Infinite 200 Pro plate reader, maintained at 37 °C. Fluorescence was recorded every 2 minutes for 2 hours using the following settings: excitation at 380 nm, emission at 650 nm, gain 220, 25 flashes, 30 µs integration delay, and 100 µs integration time.

Data were analyzed by determining the slope of the fluorescence lifetime signal (indicative of oxygen consumption) after an equilibration period of 20 minutes. The linear portion of the curve (typically over a 60-minute window) was used for slope determination. Background signal from cell-free wells was subtracted from all conditions to obtain net oxygen consumption rates. Data was normalized to the untreated control.

### Thin section electron microscopy

The *T. gondii in vitro* cyst culture was performed as described above. After four weeks, cells were treated with 5 µM atovaquone, 500 nM buparvaquone, 5 µM MMV1028806, corresponding to the lowest lethal concentration, or 0.1% DMSO, as a solvent control, for 24h. Cells were fixed by covering the monolayer with a fixation solution of 1% paraformaldehyde and 2.5% glutaraldehyde (Electron Microscopy Services) in 0.05 M HEPES buffer (pH 7.4). After incubation at RT for 3h, samples were sealed with parafilm and stored in the fridge until further processing. Fixed cells were scraped, sedimented by centrifugation (3000 *g*, 10 min) using a swing-out rotor and washed twice with 0.05 M Hepes buffer. The cell sediment was embedded in low melting-point agarose and stored in 2.5% glutaraldehyde in 0.05 M HEPES buffer. Small blocks of the embedded sediment were subjected to postfixation (osmium tetroxide, tannic acid), *en bloc* contrasting (uranyl acetate), dehydration, and embedding in epoxy resin according to a standard protocol (Laue 2010).

Ultrathin sections were prepared with an ultramicrotome (UC7, Leica Microsystems, Germany) using a diamond knife (45°, Diatome, Switzerland). The sections were collected on bare copper grids (300 mesh, hexagonal mesh shape) or copper slot grids, contrasted with 2% uranyl acetate and 0.1% lead citrate, and coated with a thin (2-3 nm) layer of carbon. The ultrathin sections with a thickness of ∼65 nm were examined with a transmission electron microscope (Tecnai Spirit, Thermo Fisher Scientific) operated at 120 kV. Images were recorded with a side-mounted CCD camera (Phurona, EMSIS, Germany) at 4095 x 2984 pixels.

### Metabolomic analysis of drug-treated intracellular tachyzoites and bradyzoites

HFF cells were infected with PruTom at MOI 1. Approximately 64 h later, the cultures were treated with 1.5 µM MMV019189, 7 µM MMV1028806, 30 µM MMV228911, 1.2 µM MMV595321, 5 µM MMV688762, 2 µM MMV688854, respectively, and 0.1 % DMSO as a control for 3h. These concentrations correspond to the five-fold respective IC_50_ value. Cultures were quenched in ice-cold PBS and parasites were isolated from their host cells by passaging scraped monolayers through a 23G needle, filtered through a 3 µm polycarbonate filter and after microscopic counting distributed at 1 x 10^8^ parasites per sample. Parasites were washed with ice-cold PBS via 5 min centrifugation at 21,500 x *g* at 0 °C. Metabolites were then extracted by sonication for 2 min in 50 µL 80 % acetonitrile in water. The insoluble particles were pelleted for 5 min at 21,500 x *g* at 0 °C. 5 µL of each sample was collected to generate a pooled biological quality control (PBQC) which was injected regularly within the otherwise randomized run list. 5 µL of each sample was injected onto either an ACQUITY UPLC BEH amide column (Waters) equipped with a VanGuard amide pre-column (Waters) or onto a SeQuant ZIC-pHILIC (Merck) column with an OPTI-LYNX ZIC-pHILIC guard column cartridge (Optimize Technologies) operating on a Vanquish Flex (Thermo Fisher). Analytes were chromatographically separated starting with 100 % eluent A (90 % acetonitrile in water with 10 mM ammonium carbonate) and 0 % eluent B (10 mM ammonium carbonate in water), decreasing to 40% A and 60% B over 25 min. Metabolites were detected on a Q Exactive Plus (Thermo Fisher) equipped with an electrospray ionization source operated in rapid polarity-switching mode at an MS^1^ resolution of 70k and an MS^2^ resolution of 35k (Top3 MSX1). Peak extraction, retention time alignment, and peak annotation were computed with Compound Discoverer 3.1 (Thermo Fisher) using an in-house library of authentic standards. Blank subtraction, gap-filling as well as statistical analysis were performed in Excel (Microsoft), and the data was plotted with GraphPad Prism 8.4.1. Principal component analysis was performed with ClustVis (Metsalu and Vilo 2015).

### Isotopic labeling

Intracellular parasites were treated as mentioned above and incubated for three hours with DMEM containing either 25 mM U-^13^C-glucose or 4 mM ^15^N-amide-glutamine instead of the respective unlabeled carbon sources. Quenching, parasite isolation, metabolite extraction and liquid chromatography were performed as described. To increase coverage of the isotopologues, metabolites were measured in positive and negative mode separately without MS^2^ scans but with an increased MS^1^ scan resolution of 140k. For peak identification purposes, the unlabeled DMSO-treated sample was analyzed with intermittent MS^2^ scans as described above.

### ATP luciferase assay

PruTom tachyzoites were cultivated in HFF cells without glucose, isolated 48 h post-infection and treated with 1 µM atovaquone or HDQ for 3h. PruTom bradyzoites were matured for three weeks, glucose-starved for an additional week and treated for 24 h with 1 µM atovaquone or HDQ before being released by needle passage 23G, pepsin digestion for 20 min at 37 °C. The digestion reaction was stopped by neutralizing in 1.2 % sodium bicarbonate (pH ≈ 8.3) with 0.0159 g/l phenol red as pH indicator dye (Christiansen et al. 2022). They were isolated by douncing on ice and filtered through a 3 µm polycarbonate filter. 10^5^ parasites were then aliquoted in 100 µL ice-cold PBS in a 96-well plate. ATP was quantified using BacTiter-Glo™ Microbial Cell Viability Assay (Promega) according to the manufacturer’s protocol using a standard curve. Data were plotted in GraphPad Prism 8.4.1..

### Metabolomic analysis drug responses on *in vitro* cysts

Intracellular cysts were isolated from their myotube host cells and extracted as described in (Christiansen et al. 2022; Maus et al. 2024). In short, cysts were released by needle passage (23G) and captured with DBA-coated magnetic beads. PBS (with 2% BSA) washed cysts were extracted in 80% acetonitrile and analyzed as described above. One dish yielded approximately 10^6^ cysts and was distributed into 3 samples. Prior to a statistical analysis, the dataset was normalized by dividing the compound intensities through the summed intensities of the respective samples.

## Results

### The Pathogen Box contains dually active compounds

To initially identify compounds that simultaneously target the tachyzoite and the bradyzoite stage of *T. gondii*, we tested 371 of 400 compounds from the Pathogen Box at a concentration of 10 µM against both stages. We excluded previously reported active compounds against *T. gondii* tachyzoites (Spalenka et al. 2018) from this initial screen. HFF cells were infected with tachyzoites of the Pru strain expressing tandem Tomato fluorescent protein in its cytosol (PruTom) at an MOI of 1 (Fig. 1A). 47 compounds completely inhibited growth during 7 days of exposure while the parasitemia of DMSO-treated controls already peaked after 4 days (Fig. 1B). To also identify bradyzocidal compounds, we differentiated KD3 myoblasts for one week into multinucleated myotubes and infected them under bicarbonate-deplete conditions and at a multiplicity of infection of 0.1 for four weeks to allow maturation of tissue cysts (Fig. S1). We exposed the cultures to 10 µM of the 371 MMV compounds for one week, removed the inhibitors and stimulated re-differentiation into tachyzoites by bicarbonate supplementation (Fig. 1C). Growth of DMSO-treated cultures became detectable after 10 to 14 days. Interestingly, 31 compounds prevented any tachyzoite regrowth (Fig. 1D), indicating their bradyzocidal activity. To exclude host cytotoxicity-dependent suppression of parasite growth, we measured cell viability via the reductive potential of myotubes and HFF cells using a resazurin assay. At 10 µM, 30, 12 and 13 compounds inflicted cytotoxicity onto HFFs, myotubes or both cell types, respectively (Supplemental File 1). This initial screen identified 14 compounds with activity against both tachyzoites and bradyzoites and no apparent cytotoxicity (Fig. 1E).

### Validation of screening hits

To validate our results, we determined half-inhibitory concentrations (IC_50_) in PruTom tachyzoites (Fig. 2A, B) and half-maximal lethal concentration (LC_50_) in bradyzoites. We also included 29 compounds that were previously reported to target tachyzoites (Spalenka et al. 2018). We confirmed dual activity of 16 compounds that exhibited mean IC_50_ values ranging between 70 pM and 1.1 µM for tachyzoites (Fig. 2E, Fig. S2D) and LC_50_ values between 0.15 and 8.2 µM for bradyzoites (Fig. 2F, Fig. S2D). To allow a comparison between the tachyzoite and bradyzoite data, we also determined the respective minimal lethal concentrations (MLC) over a seven-day treatment duration of tachyzoites as well (Fig. 2G). Except for four not significantly different compounds (MMV021013, MMV022478, MMV658988, MMV65900), minimal lethal concentrations appear generally higher in bradyzoites, potentially reflecting their lower metabolic activity and/or the cyst wall as a diffusion barrier.

### Dually active MMV Pathogen Box hits share high lipophilicity and are indicated to be active against other pathogens

To gain insight into general properties of active compounds from our initial screen (Fig. 1E), we tabulated indicated activities as provided by MMV (Fig. S2A, D). As expected, these compounds were mostly earmarked as active against *T. gondii* tachyzoites (67%), however, none of those compounds exhibited bradyzocidal activity. Instead, 27% of dually active compounds were considered active against *Cryptosporidium*. Also, despite the relative phylogenetic proximity of *P. falciparum,* only 5% of the 125 potential antimalarials blocked growth of both *T. gondii* stages. To characterize physicochemical properties of these dually active hits, we approximated their lipophilicity by their logP values (Supplemental File 1) and compared them to compounds that were inactive against *T. gondii* (Fig. S2B). We found that tachyzocidal, bradyzocidal and dually active compounds possess a statistically significantly higher lipophilicity than non-active compounds and this trend appeared more accentuated for bradyzocidal and dually active compounds. However, when comparing tachyzocidal with bradyzocidal compounds, including dually active ones, we did not find a difference (p = 0.06). We estimated permeability through the blood brain barrier (bbb) using LightBBB (Shaker et al. 2020) and gastrointestinal absorbability using SwissADME (Daina, Michielin, and Zoete 2017) (Supplemental File 1). Most of the dually active drugs were predicted to be both permeable across the bbb and absorbed in the intestine (Fig. S2C). Together, these data support phylogenetic relationship as a weak predictor of activity of Pathogen Box compounds against *T. gondii* bradyzoites and highlight the role of lipophilicity among the compounds tested here.

### Metabolic drug responses in tachyzoites indicate conserved and novel modes of action

We chose to focus on dually active compounds, because they are clinically more relevant and investigation of their targets allows comparisons between on parasite stages. To that end, we choose five compounds based on their availability and yet unknown mode of action (Fig. 3, Supplemental File 2, Fig S3C) to test metabolite responses of tachyzoites. Infected monolayers were treated for three hours with compound concentrations five times their respective IC₅₀ values or the solvent DMSO. The inhibitor MMV688854 (BKI 1294) (Doggett et al. 2014) is considered to inhibit a non-metabolic target (calcium-dependent protein kinase (CDPK1)) at the employed concentration (Hayward et al. 2023) and served as an additional control to estimate the metabolic impact of growth inhibition. We filter-purified parasites from quenched monolayers and analyzed their acetonitrile extracts by LCMS using both, pHILIC and BEH-amide chromatography columns. We generated 9 samples per condition from three repeats of the experiment with triplicate samples. A principal component analysis illustrates that batch effects are the main contributors to biological variances within the dataset (Fig. S3A, B). Despite that, all compounds elicited a distinct metabolic phenotype on both LCMS systems (Fig. 3). The CDPK1 inhibitor increased levels of AMP, oxidized glutathione, pseudo uridine, citrulline, glycolate and glutamyl glutamine, while reducing the abundances of GABA, carbamoyl aspartate, phosphoenolpyruvate and a hexose phosphate. Shifts in these metabolites are shared among many treatments and indicate their implication in a general stress response and thus do not signify direct interference with parasite metabolism.

**Figure 3:**
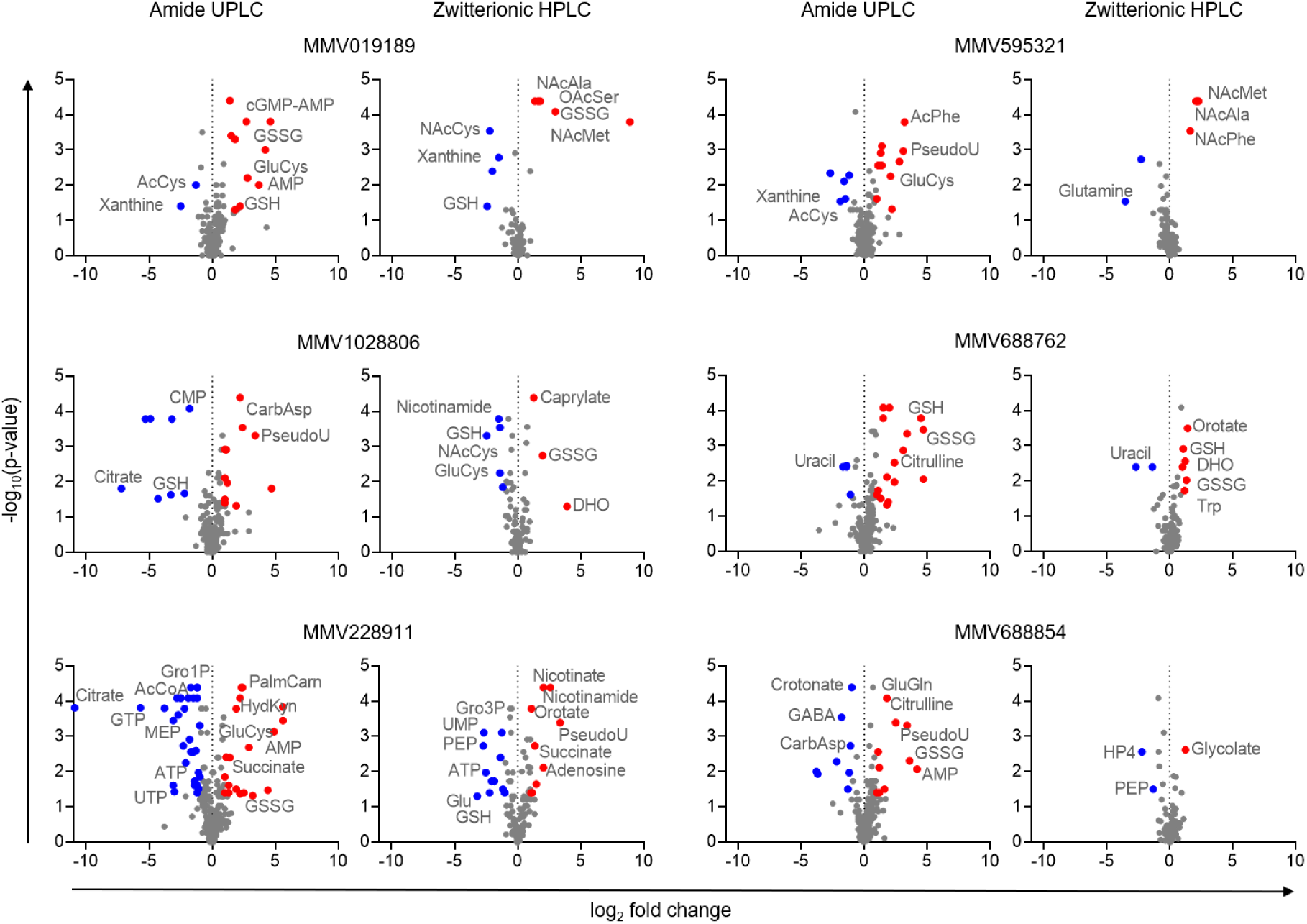
Untargeted metabolomics of tachyzoites treated with 6 screening hits effective against both stages of *T. gondii*. Volcano plots showing the parasitic, intracellular metabolic phenotype caused by MMV Pathogen Box compounds that inhibited tachyzoites and bradyzoites during the screen. The metabolome was measured with two columns of complementary chemistries to enhance metabolite coverage. Blue dots indicate metabolites which were significantly reduced compared to DMSO treated parasites and red dots indicate significantly accumulated metabolites (p < 0.05, log2 fold change <-1 or > 1, 3x biological replicates with 3x technical replicates each, n = 9). Abbreviations: AcCoA - Acetyl-CoA, AcCys - Acetylcysteine, AcPhe - Acetylphenylalanine, AMP - Adenosinemonophosphate, ATP - Adenosinetriphosphate, CarbAsp - Carbamoylaspartate, cGMP-AMP - cyclic Guanosinemonophosphate-adenosinemonophosphate, CMP - Cytosinemonophosphate, DHO - Dihydroorotate, GABA - gamma-Aminobutyratic acid, GluCys - Glutamylcysteine, GluGln - Glutamylglutamine, Gro1P – Glyceraldehyde 1-phosphate, Gro3P – Glyceraldehyde 3-phosphate, GSH – Glutathione oxidized, GSSG – Glutathione reduced, GTP – Guanosinetriphosphate, HP4 – Hexose phosphate No 4 (fourth chromatographic peak of 260.02972), HydKyn – Hydroxylkynurenine, MEP – 2-C-Methyl-D-erythritol 4-phosphate, NacAla – *N*-Acetylalanine, NacCys – N-Acetylcysteine, NacMet – *N*-Acetylmethionine, NacPhe – *N*-Acetylphenylalanine, OacSer – *O*-Acetylserine, PalmCarn – Palmitoylcarnitine, PEP – Phosphoenolpyruvate, PseudoU – Pseudouridine, Trp – Tryptophan, UMP – Uridinemonophosphate, UTP – Uridinetriphosphate

Inhibitor-specific responses include the accumulation of cGMP-AMP after treatment with MMV019189, accumulation of carbamoyl aspartate and dihydroorotate upon MMV1028806 treatment, and a general depletion of central carbon metabolites (succinate, citrate, glyceraldehyde 3-phosphate, ATP, MEP) in MMV228911-treated parasites. MMV595321 caused an accumulation of N-acetylated amino acids, and MMV688762 led to uracil deficiency and an accumulation of dihydroorotate and orotate. These data suggest that growth inhibitors cause a bipartite metabolic response composed of generic and compound-specific changes. While these findings highlight compound-specific metabolic signatures, some observed alterations may represent downstream or systemic effects of inhibitor action, reflecting the inherent complexity of parasite metabolism.

### MMV1028806 and buparvaquone elicit metabolic responses similar to the *bc1*-complex inhibitor atovaquone

The metabolic impact of most tested MMV compounds indicated a novel mode of action that did not resemble known or recognizable metabolic fingerprints. However, the metabolic fingerprint of MMV1028806 was consistent with inhibition of pyrimidine biosynthesis at the *bc*_1_-complex-dependent oxidation reaction of dihydroorotate to orotate, as has been reported for *P. falciparum* (Fig. 4A) (Allman et al. 2016; Creek et al. 2016; Antonova-Koch et al. 2018). A similar response would be expected to buparvaquone, which we found to be dually active against *T. gondii* tachyzoites and bradyzoites and which targets the *bc*_1_-complex in *Theileria annulata* (McHardy et al. 1985), *Neospora caninum* (Müller et al. 2015) and *T. gondii* tachyzoites (Hayward et al. 2023).

**Figure 4:**
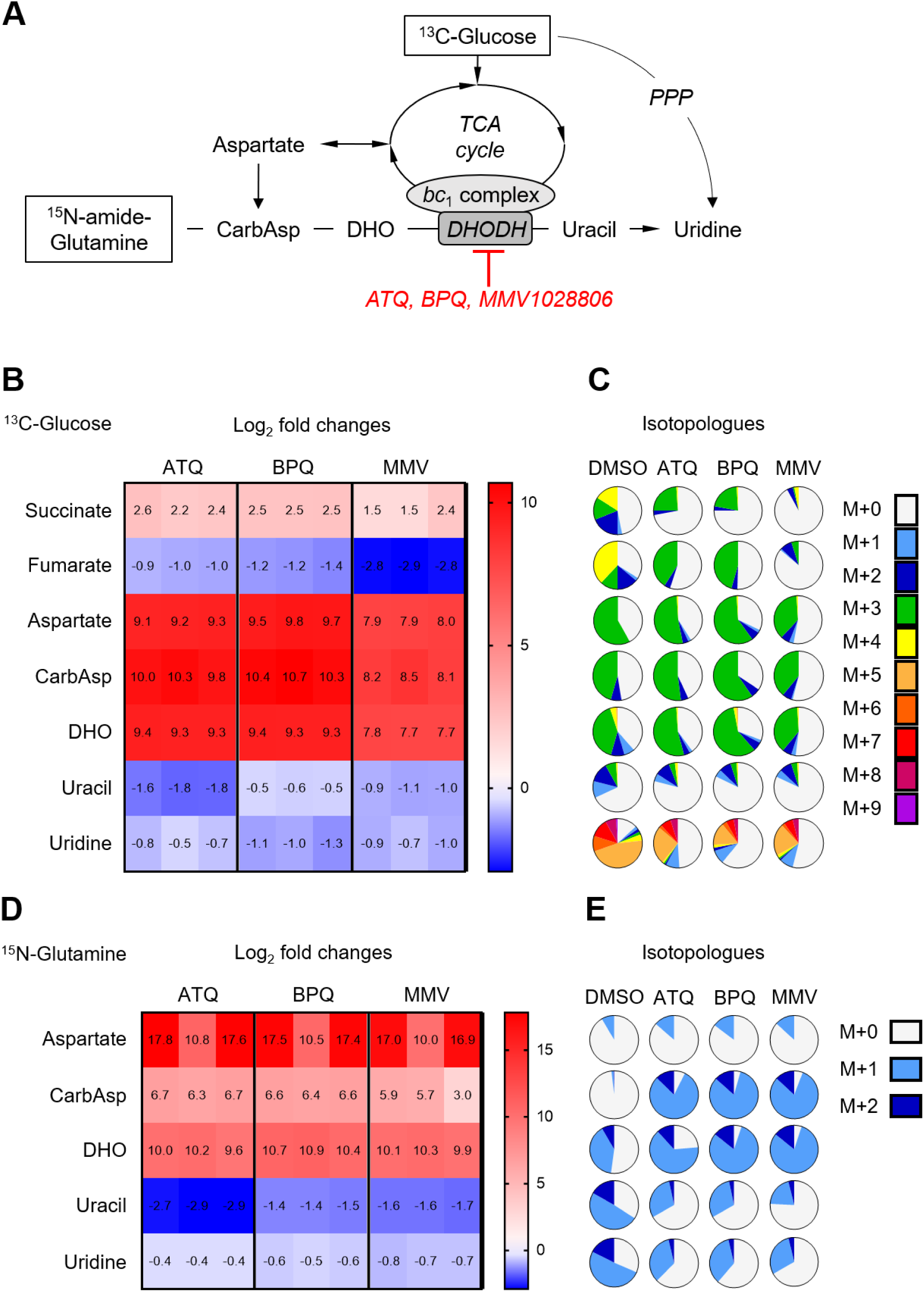
The metabolic response to electron transport chain inhibitors and MMV1028806. (A) Scheme of the affected pathways upon *bc*_1_-complex inhibition. Intracellular tachyzoites were grown in presence of U-^13^C-glucose (B, C) or ^15^N-amide-glutamine (D, E) instead of unlabeled carbon sources and treated with atovaquone (ATQ), buparvaquone (BPQ) and MMV1028806 (MMV) for 3h. (B, D) The treatment induced changes of mETC-related metabolite abundances are shown as log_2_-fold changes in comparison to DMSO treated cultures. Shown are three replicate measurements. (C, E) Shown is the average isotopologue distribution of mETC-related metabolites. The pie charts depict the number of heavy carbon or nitrogen atoms incorporated into the respective metabolites (M+0 in light grey represents the unlabeled metabolite, while M+1 in light blue indicates the integration of one heavy atom into the molecule). Shown are the means of three replicate measurements. Abbreviations: ATQ – atovaquone, BPQ – buparvaquone, CarbAsp – carbamoyl-aspartate, DHO – dihydroorotate, DHODH – dihydroorotate dehydrogenase, GABA – gamma-aminobutyrate, MMV – MMV1028806, PPP – pentose phosphate pathway, SSA – succinic semialdehyde

To test these hypotheses, we exposed intracellular parasites to MMV1028806 or BPQ while substituting glucose with ^13^C6-glucose for three hours. We then measured the incorporation of ^13^C into the central carbon metabolites as well as the intermediates of the pyrimidine synthesis pathway. As controls, we utilized the known *bc*_1_-complex inhibitor atovaquone and the solvent DMSO (Supplemental File 3). Treatment with all three inhibitors led to a dramatic accumulation of +3 labeled aspartic acid, carbamoyl aspartate and dihydroorotate which was accompanied by a depletion of uracil and uridine, in particular of the +5-labeled uridine isotopologue. These changes are consistent with impaired function of the *bc*_1_-complex-dependent dihydroorotate dehydrogenase (Fig. 4 B, C). Another common effect was the accumulation of succinate and depletion of fumarate, suggesting reduced recycling of FADH_2_ via the *bc*_1_-complex.

To independently verify our results, we hypothesized that *T. gondii* also incorporates glutamine-derived nitrogen into pyrimidines. We replaced glutamine with its ^15^N-amide-labeled isotopologue in the tachyzoite culture media for three hours while simultaneously exposing the parasites to ATQ, BPQ or MMV1028806. As expected from our previous experiments, we observed a marked decrease of uracil and uridine levels while aspartate, carbamoyl aspartate and dihydroorotate accumulated again (Fig. 4D, Supplementary File 5). ^15^N was incorporated into many metabolites (Fig. S4). Consistently, labelling of pyrimidines was diminished while carbamoyl aspartate and dihydroorotate were heavily labeled as M+1 isotopologues (Fig. 4E).

Together, these data show that treatments with all three compounds lead to the accumulation of metabolites upstream of the *bc*_1_-complex-dependent reactions during pyrimidine biosynthesis and the TCA cycle and to the depletion of downstream metabolites (Fig. 4A). These metabolic signatures suggest an inhibition of the *T. gondii bc*_1_-complex in tachyzoites.

### Atovaquone, buparvaquone and MMV1028806 decrease the mitochondrial membrane potential of tachyzoites and bradyzoites and block the electron transport chain

The inhibition of the *bc*_1_-complex by ATQ leads to the collapse of the mitochondrial membrane potential in *T. gondii* tachyzoites (Vercesi et al. 1998) and we anticipated a similar phenotype in BPQ or MMV1028806-treated parasites. We treated intracellular tachyzoites and bradyzoites for 24 h with the three inhibitors or 0.1 % DMSO alone and estimated the mitochondrial membrane potential and morphology using a fluorescent Mitotracker dye. The mitochondria of control tachyzoites appeared as extended loops but largely collapsed after treatments with all three compounds (Fig. 5A, B). Notably, also host mitochondria appear influenced. To evaluate the specificity of this assay we used pyrimethamine, clindamycin and 6-diazo-5-oxonorleucine (DON) that target dihydrofolate reductase, apicoplast translation and glutaminase as controls. Mitotracker and GFP signal intensities remained comparable to DMSO controls (Fig. S5). DAPI signal intensity appeared reduced in pyrimethamine-treated samples. In bradyzoites, mitochondria appeared mostly as spheres or were donut-shaped (Supplementary File 6) and their relative fluorescence decreased after 24 h treatment with ATQ, BPQ or MMV1028806 at 5 µM, 500 nM or 5 µM, respectively while their morphology appeared unchanged (Fig. 5C, D). Analysis by thin section electron microscopy revealed a largely unaffected sub-mitochondrial ultrastructure but the areas of mitochondrial profiles were changed in comparison to control after exposure with ATQ and MMV1028806 but not with BPQ (Fig. S6). This indicates that the latter decreases Mitotracker fluorescence intensity directly, while ATQ and MMV1028806 cause both, a drop in Mitotracker fluorescence and a reduced size of the mitochondria which leaves open whether a change of the membrane potential, of the size, or even of both parameters were responsible for the drop in Mitotracker fluorescence.

**Figure 5:**
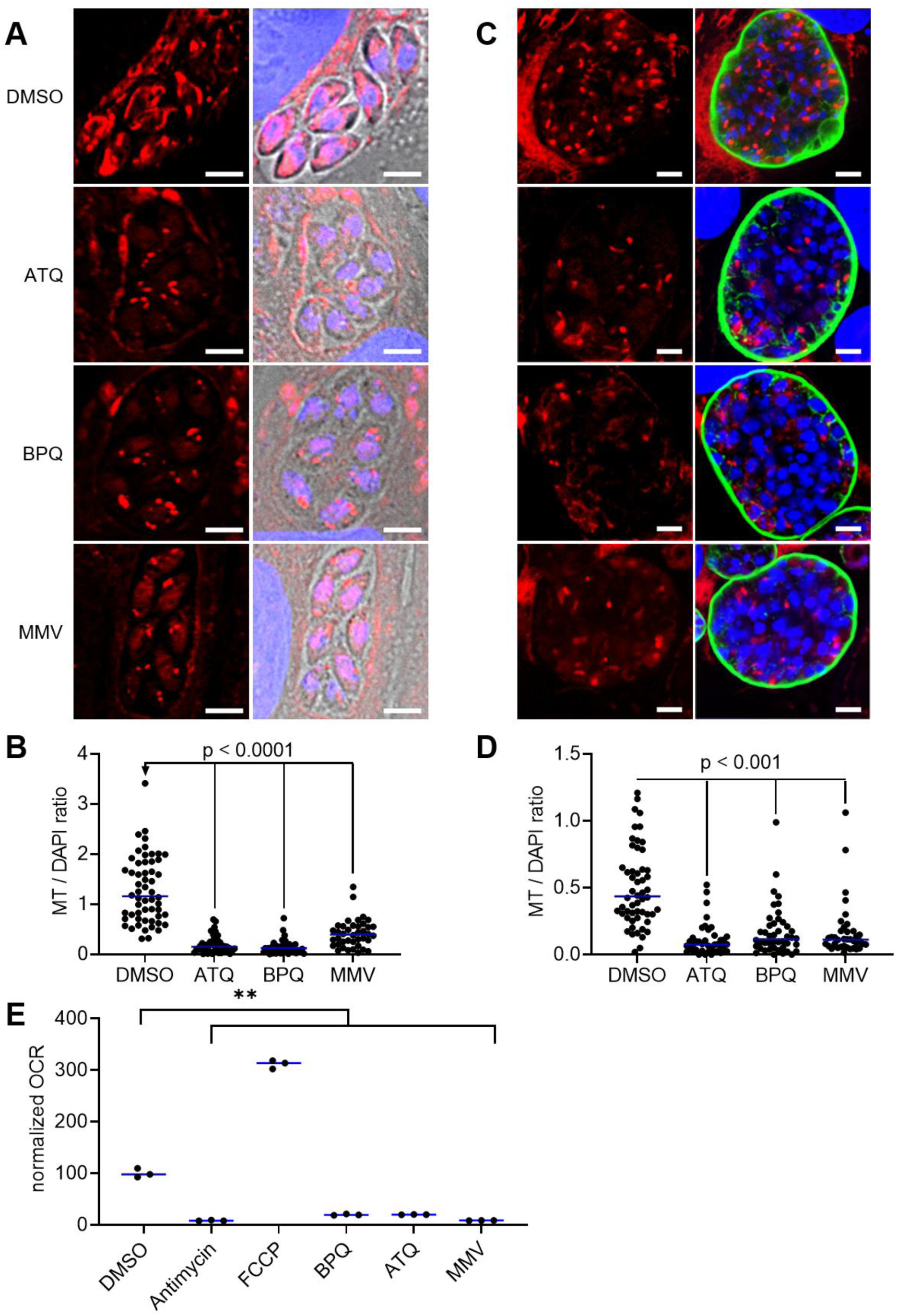
Atovaquone, buparvaquone, and MMV1028806 reduce mitochondrial potential and respiration in *T. gondii*. (A) HFF cells were infected with tachyzoites and treated with DMSO, atovaquone, buparvaquone and MMV1028806 for 24 h, and mitochondria were stained with Mitotracker (red) and DNA was stained with DAPI (blue). scale bar = 5 µm. (B) The ratios of fluorescence intensities between Mitotracker (MT) and DAPI dyes were calculated per vacuole (DMSO n = 56, ATQ n = 45, BPQ n = 49, MMV1028806 n = 40; Blue lines represent the median, two-sided Mann-Whitney test). (C) 4 weeks old, myotube derived ME49 in vitro cysts were treated with DMSO, atovaquone, buparvaquone and MMV1028806 for 24 h, and mitochondria were stained with Mitotracker (red), the cyst wall was stained with Dolichos Biflorus Agglutinin and Streptavidin-Cy2 (green), and DNA was stained with DAPI (blue). scale bar = 20 µm. (D) The ratios of fluorescence intensities between Mitotracker (MT) and DAPI dyes were calculated per vacuole (DMSO n = 56, ATQ n = 42, BPQ n = 45, MMV1028806 n = 38; Blue lines represent the median, two-sided Mann-Whitney test). (E) 10 million freshly egressed tachyzoites were incubated with known *bc*_1_ complex inhibitors ATQ and BPQ, specific control compounds Antimycin and FCCP, and MMV1028806. Every treatment significantly impacts the oxygen consumption rate (OCR) (blue line represents the median, n = 3) (Welch’s test; ** p < 0.005).

We next tested directly whether the mETC was blocked by ATQ, BPQ and MMV1028806 by measuring oxygen consumption of treated tachyzoites using a fluorescence-based assay (Fig. 5E). The solvent DMSO, the decoupling agent carbonyl cyanide 4-(trifluoromethoxy)phenylhydrazone (FCCP), and the *bc1*-complex inhibitor antimycin A served as controls. Treatment with FCCP led to a marked increase in oxygen consumption compared to the DMSO control, indicating maximal respiratory capacity of the parasites. In contrast, treatments with ATQ, BPQ, MMV1028806, and antimycin A resulted in substantially reduced oxygen consumption levels relative to the DMSO control and suggest indeed a blockage of the mETC consistent with the inhibition of the *bc1*-complex.

### Inhibitors of the mitochondrial electron transport chain alter catabolic processes in bradyzoites

Next, we sought to confirm this mode of action directly on bradyzoites and record the metabolic impact of ATQ, BPQ or MMV1028806. To that end, we exposed intracellular four-week-old cysts to 5 µM, 500 nM or 5 µM of the respective compounds and for three hours and used 0.2% DMSO as a mock treatment. These concentrations represent their LC_50_ against bradyzoites and were chosen to minimize off-target effects. The cysts were isolated for LCMS analysis using magnetic beads (Christiansen et al. 2022) (Fig. 6A). Common in all three distinct metabolic phenotypes (Fig. S7) is the accumulation of AMP and NADH, while varying TCA cycle intermediates, such as malate, succinate and α-ketoglutarate and mitochondria-independent energy sources such as glucose and phosphocreatine were depleted (Fig. 6B-D). These metabolic changes suggest an energy starvation scenario marked by rising AMP levels and reduced recycling of NADH via the *bc*_1_-complex. Interestingly, in BPQ and MMV1028806 treated cysts, we also observed prominent accumulation of several acyl-carnitines species which are mobile intermediates of ß-oxidation (Fig. 6B-D). Since this pathway is thought to be absent in *T. gondii,* we attribute these changes to potential inhibitory effects on host mitochondria. Notably, we did not observe changes in the pyrimidine synthesis pathway, which may reflect low nucleic acid demand from largely dormant bradyzoites. These data demonstrate that bradyzoites, which were considered metabolically inactive, exhibit metabolic responses to inhibitory compounds. Although we characterized AMP accumulation as part of a general stress response in tachyzoites, the metabolic profiles of inhibited bradyzoites are consistent with inhibition of their mitochondrial *bc*_1_-complex and suggest its role in ATP generation.

**Figure 6:**
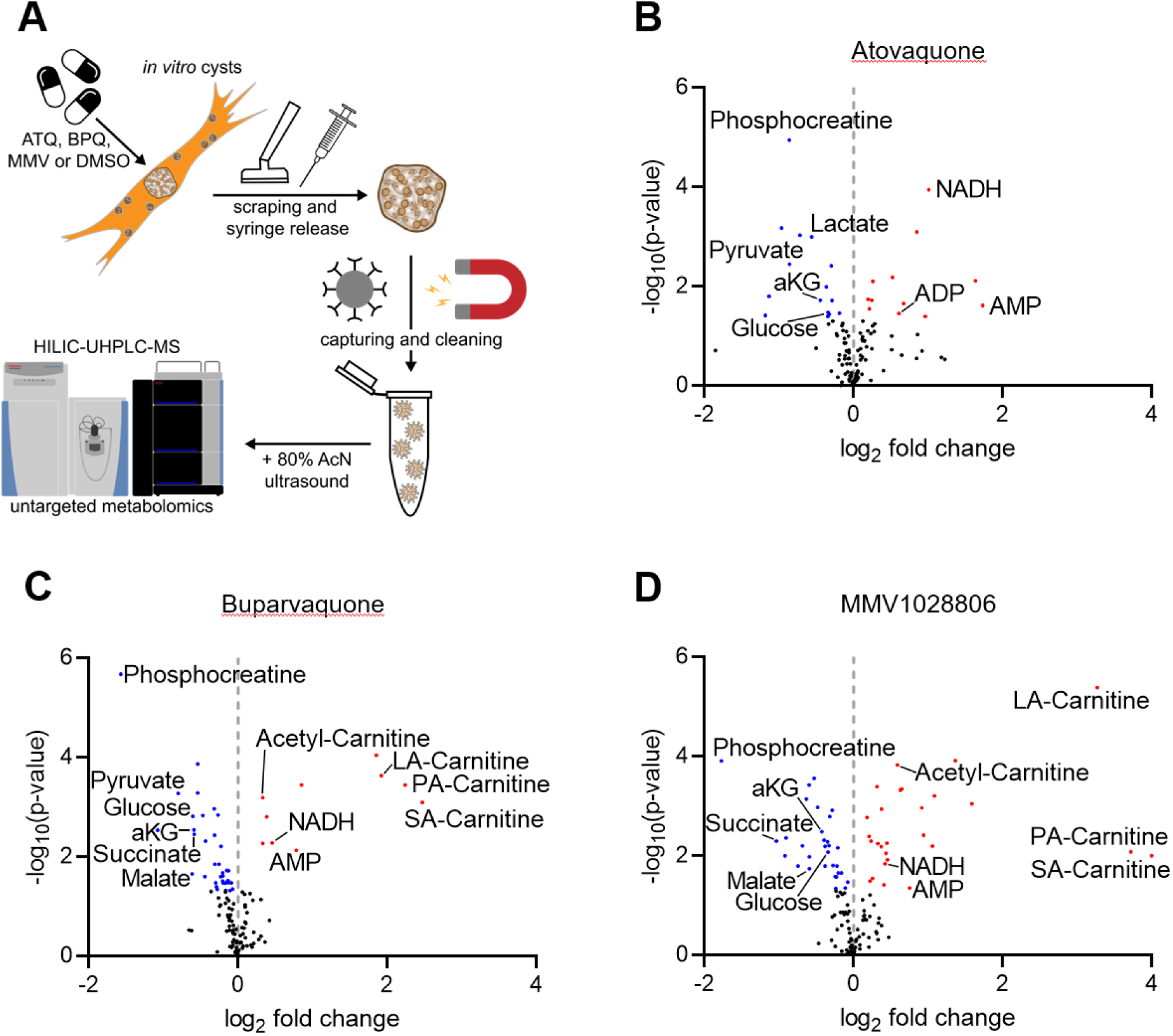
Untargeted metabolomic analysis of bradyzoites treated with *bc*1-complex inhibitors shows an energy imbalance. (A) Four weeks old *in vitro* cysts were treated for three hours. The cell monolayer was scraped off and the cysts were syringe-released. With *Dolichos biflorus* agglutinin (DBA) coated magnetic beads, the isolated cysts were captured and washed with PBS. The metabolites were extracted with 80% acetonitrile (AcN) in water and sonication, and analyzed via hydrophilic-interaction ultra-high pressure liquid chromatography coupled mass spectrometry (HILIC-UHPLC-MS). The results are shown in three separate volcano plots: Metabolic phenotypes of bradyzoites treated with Atovaquone (B), Buparvaquone (C), and MMV1028806 (D). (n = 3, significance analyzed with a two-sided U-Mann-Whitney test) Abbreviations: ATQ - atovaquone, BPQ - buparvaquone, DMSO – dimethyl sulfoxide, LA-carnitine - linoleoylcarnitine, MMV - MMV1028806, NADH - reduced nicotinamide adenine dinucleotide, PA-carnitine - palmitoylcarnitine, SA-carnitine - stearoylcarnitine

### The differential efficacy of atovaquone and HDQ against bradyzoites correlates with the inhibition of ATP levels in bradyzoites

We sought to test the importance of the mETC chain in ATP generation in bradyzoites and exploited the differential targets of the coenzyme Q analogs ATQ and 1-hydroxy-2-dodecyl-4(1)quinolone (HDQ). However, while ATQ inhibits the *bc*_1_-complex, HDQ inhibits dihydroorotate dehydrogenase (DHODH) (Hegewald, Gross, and Bohne 2013) and the alternative NADH dehydrogenase (NDH2) (Saleh et al. 2007). Both compounds block tachyzoite growth at low nanomolar concentrations in vitro with IC50s at 20 and 4 nM, respectively (Fig. 7A). However, in contrast to ATQ, HDQ does not affect the viability of mature *T. gondii* bradyzoites despite diminishing their mitochondrial membrane potential (Christiansen et al. 2022). Up to 20 µM HDQ did not arrest re-differentiation into proliferating tachyzoites, while bradyzoites that were treated with 5 µM ATQ (LC_50_ 2.5 µM) did not generate proliferating tachyzoites (Fig. 7B). To investigate how these modes of actions affect the mETC in bradyzoites, we tested their influence on mitochondrial ATP production by luciferase-based ATP assay. This assay shows a linear response to parasite cell equivalents between 10^3^ and 10^6^ parasites per 100 μL, and in order to minimize variance we decided to work with 10^5^ parasites (Fig. S8). To maximize mitochondrial ATP production, we starved three-week-old bradyzoites for one additional week of glucose. As a comparison, we utilized tachyzoites that were cultured for two days in glucose-deplete conditions. Untreated tachyzoite samples contained in average 196 ± 21 fmol ATP per 10^5^ parasites, whereas untreated bradyzoites only contained 14.1 ± 1.1 fmol. (Fig. 7C). Three hour-long treatment with 5 µM ATQ decreased ATP levels in both stages by 75 % (Fig. 7D). In contrast, HDQ reduced ATP levels by 60 % in tachyzoites but only marginally affected bradyzoites (Fig. 7D). These data indicate that bradyzoites can maintain very low ATP levels and suggest the presence of a diminished but obligate production of ATP in the mitochondria that functions independently of exogenous glucose and HDQ-target enzymes.

**Figure 7:**
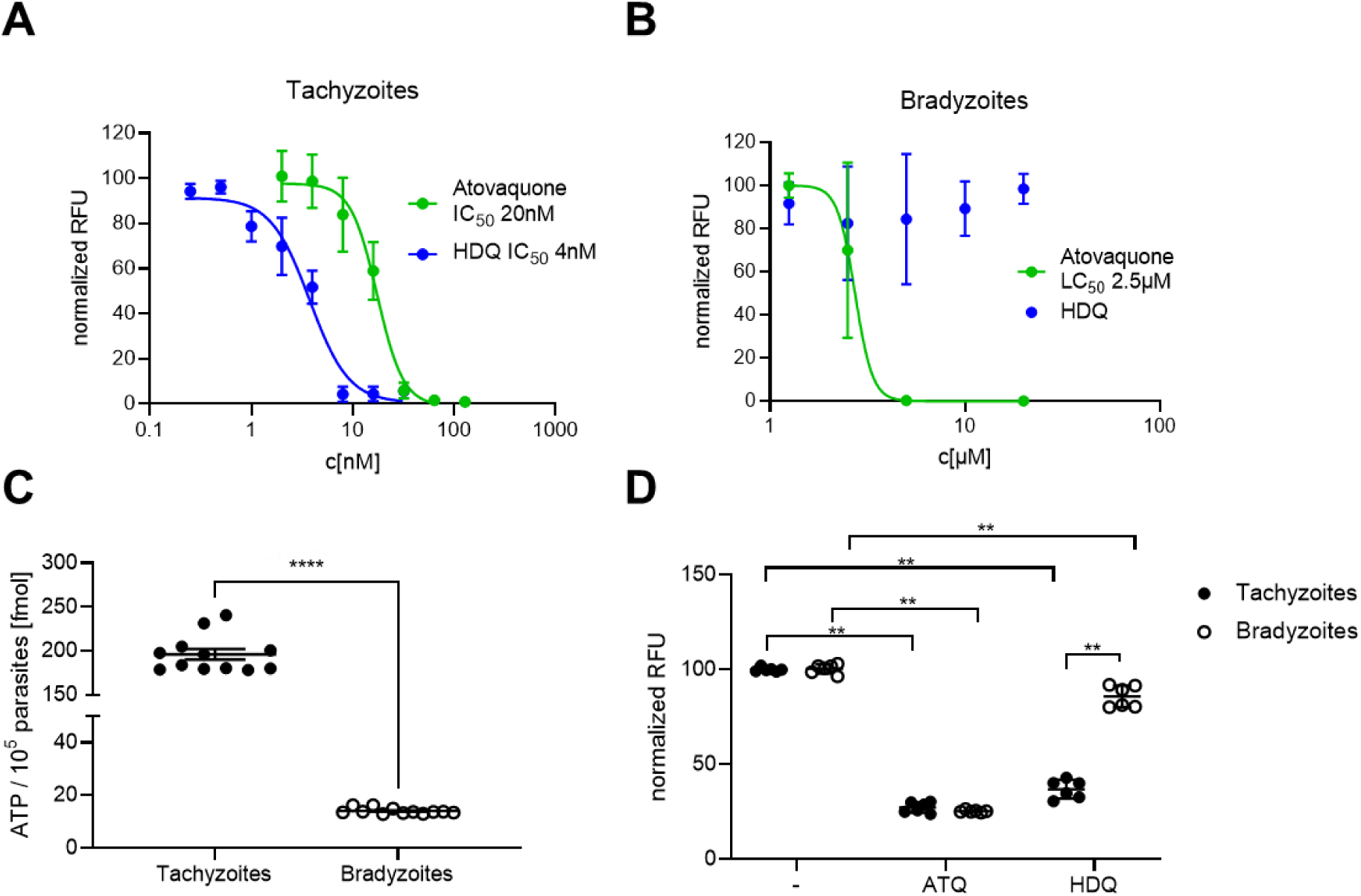
Differential efficacy and ATP depletion by atovaquone and HDQ in tachyzoites and bradyzoites. Atovaquone and HDQ were tested against *T. gondii* Pru parasites. (A) Showing the normalized half-maximal inhibitory concentration (IC_50_) against tachyzoites in fibroblasts at day 4, n=8. (B) 4-week-old bradyzoites were treated for a week with Atovaquone or HDQ. Cultures were observed for tachyzoite re-growth for three weeks. The half-maximal lethal concentration (LC_50_) was determined at the time point which ideally reflected the dose-response curve, in this case day 14, n=6. 10^5^ tachyzoites and bradyzoites were lysed and the ATP concentration was determined with an enzymatic luciferase assay. (C) Showing the calculated concentration of ATP within an average parasite. Lines reflect the median, error bars represent SEM, significance test Mann-Whitney, n=12. (D) Parasites were treated with 1μM Atovaquone or HDQ. RFUs were normalized to the untreated control of either the tachyzoite or bradyzoite dataset. Shown are means, error bars represent SD, significance test is pairwise Mann-Whitney, n=6.

## Discussion

New treatment options for chronic *T. gondii* infections are needed, as chronic infections are currently incurable and available treatments for acute toxoplasmosis are associated with severe side-effects, in particular in infants after congenital transmission (Konstantinovic et al. 2019). To simultaneously identify drug targets in bradyzoites and along with their inhibitory compounds, we utilized a recently developed *in vitro* culture system of pan-drug tolerant bradyzoites (Christiansen et al. 2022) to screen 400 compounds of the MMV Pathogen Box.

16 compounds exhibited confirmed cidal activity against bradyzoites while also inhibiting growth of tachyzoites. Metabolic profiling and stable isotope tracing in treated tachyzoites suggested the inhibition of the mitochondrial *bc_1_-*complex by MMV1028806 and the reference compound BPQ.

Interestingly, of the 15 compounds in the Toxoplasmosis disease set, previously earmarked by MMV, only 10 were found to be active against tachyzoites and none against bradyzoites. These differences may be attributed to strain-specific susceptibilities which are known for anti-folates and atovaquone (Meneceur et al. 2008) and differences in the design of growth assays. MMV assessed activities via plaque assays that prevent media change and rely on visual inspection of growth of a low number of parasites and might overestimate inhibitory effects.

We found that bradyzocidal compounds exhibit higher predicted logP coefficients, suggesting that the parasite entry might occur via diffusion across the cyst wall and less via solute transporter proteins. A known example of this principle is the anti-malarial drug fosmidomycin, where high polarity and low import rates caused minimal uptake by and efficacy against *T. gondii* tachyzoites (Nair et al. 2012; Baumeister et al. 2011). A similar case might also be made for the antifolates, pyrimethamine, sulfadiazine, trimethroprim and sulfamethoxazole which exhibit low consensus logP values of 2.09, 0.4, 1.16 and 0.92 and which also fail to eradicate tissue cysts (Alday and Doggett 2017; Enshaeieh et al. 2021). Interestingly, MMV675968 is a dually active compound with a logP value of 2.46, that has been shown to be a potent inhibitor of the dihydrofolate reductase (DHFR) in a range of pathogens (Songsungthong et al. 2019, 2021; Kim et al. 2021; Borba-Santos, Vila, and Rozental 2020; Nugraha et al. 2019; Rollin-Pinheiro et al. 2021; Vila and Lopez-Ribot 2017; Cantillon et al. 2022). Its target in *T. gondii*, however, remains to be identified.

Another important diffusion barrier that needs to be overcome to eradicate *T. gondii* tissue cysts is the blood brain barrier (BBB). Pyrimethamine and trimethroprim penetrate the blood brain barrier (Geils et al. 1971; Ineichen et al. 2020), they do not clear brain-resident cysts (Montazeri et al. 2018; Enshaeieh et al. 2021; Christiansen et al. 2022) highlighting the cyst wall as a potentially decisive obstacle. Consistent with this hypothesis is the tolerance of mature in vitro tissue cysts against high dose of sulfadiazine and pyrimethamine (Christiansen et al. 2022).

To characterize the modes of action of four additional inhibitors, we used metabolomics. In general, metabolic phenotypes reflect direct consequences of target inhibition as well as indirect effects resulting from attenuated growth and stress responses. We chose the bumped-kinase inhibitor MMV688854 (BKI-1294) that targets the calcium-dependent protein kinase 1 (CDPK1) as a control compound to identify metabolites implicated in a general stress response and adaptation to slow growth (Winzer et al. 2015). CDPK1 is needed for invasion (Johnson et al. 2012) and also affects formation of daughter cells. An accumulation of AMP and oxidized glutathione are the most conserved responses across the six treatments and may indicate depleted energy levels and oxidative stress (Caro et al. 2012). Also, pseudo uridine is elevated in many treatments. Enhanced pseudo uridylation of RNA is considered to be implicated in stress responses (Cerneckis et al. 2022) and modulates RNA stability (Davis 1995). It remains to be tested which of these conserved metabolic responses signify cellular decay and which ones take part in a productive metabolic response in *T. gondii*. Nonetheless, metabolic profiling data on antibiotics-exposed bacterial pathogens has been used to design combination therapies, by either interfering with productive metabolic stress responses (Zampieri et al. 2017) or by simultaneously inhibiting otherwise non-essential pathways (Campos and Zampieri 2019). We found that the systematic identification of target structures and drug modes of action from profiling data alone remains challenging due to inherent non-linear relationships within the cellular metabolic network, the underlying regulatory mechanisms of enzyme functions. A systematic mapping of genetic disruptions to metabolic phenotypes, as demonstrated in *E. coli,* may alleviate these restrictions (Anglada-Girotto et al. 2022).

Apart from the general metabolic stress response to the treatment with MMV688854, three compounds caused apparent inhibitor-specific signatures. MMV019189, which is known to inhibit *P. falciparum* (Duffy et al. 2017), but also *C. parvum* (Hennessey et al. 2018) and *L. mexicana* (Berry et al. 2018), lead to the accumulation of c-GMP-AMP in the parasite. This dinucleotide is produced by the c-GMP-AMP synthase (cGAS) in mammalian cells in response to cytosolic DNA and triggers a type-I interferon response (Sun et al. 2013). While cGas is considered absent in *T. gondii*, host c-GMP-AMP induces STING-signaling as a response to *T. gondii* infection (P. Wang et al. 2019). Hence, MMV019189 may implicate the host in its antiparasitic activity.

MMV228911-treated tachyzoites showed the depletion of the DXR product 2-C-methylerythritol 4-phosphate (MEP), a metabolite that occurs in the non-mevalonate pathway in *T. gondii’*s apicoplast. Other metabolites upstream of DXR are either not detected or depleted, such as glyceraldehyde 3-phosphate, but also partake in other pathways. Interestingly, MMV228911 also caused a broader dysregulation of central carbon metabolism marked by a decrease of many phosphorylated metabolites such as nucleotide triphosphates and PEP but also citric acid and acetyl-CoA. This is consistent with metabolic signaling beyond the apicoplast, which also has been implicated in *P. falciparum*, where a cytosolic sugar phosphatase is responsible for resistance to the DXR inhibitor Fosmidomycin (Guggisberg et al. 2014).

We identified MMV1028806 as a candidate *bc*_1_-complex inhibitor by comparing its metabolic impact with the one of BPQ with ATQ. All three compounds cause a sharp increase of pyrimidine synthesis substrates, depletion of pyrimidines and dysregulation of TCA cycle intermediates. This is consistent with previous mass spectrometry-based studies that characterized metabolic fingerprint of atovaquone on *P. falciparum* intraerythrocytic blood-stages (Creek et al. 2016; Allman et al. 2016; Antonova-Koch et al. 2018). In addition to abundance data, the incorporation of ¹³C and ¹⁵N stable isotopes from glucose and glutamine, respectively, into TCA cycle and pyrimidine biosynthesis intermediates suggest the *bc1*-complex as a target. Both, the isotopologue distributions and metabolite abundances strongly suggest attenuation of pyrimidine production and TCA cycle fluxes by all three inhibitors (Hortua Triana et al. 2016). TCA cycle intermediates show characteristic isotopologue distributions that are indicative of a ^13^C-label incorporation via acetyl-CoA (MacRae et al. 2012). A relatively increased M+3 isotopologue species in dicarboxylic acids in ATQ- and BPQ-treated parasites may indicate enhanced anaplerotic synthesis from glycolytic intermediates. These distinctions from MMV1028806 metabotypes may reflect a combination of off-target effects and differential timing of the metabolic inhibition.

Interestingly, six compounds from the MMV Pathogen Box have recently been shown to inhibit O_2_ consumption in tachyzoites through *bc*_1_-complex inhibition (Hayward et al. 2023). We co-identified Buparvaquone as a *bc*_1_-complex inhibitor but we did not test the remaining five compounds for their targets. However, in our screen at 10 µM we found MMV024397 and MMV688853 to be dually active, while Trifloxystrobin (MMV688754) and Auranofin (MMV688978) lacked inhibitory activity in bradyzoites, and Azoxystrobin (MMV021057) was inactive. In addition, the previously described *bc*_1_-complex inhibitor ELQ400 (MMV671636) (McConnell et al. 2018) was dually active. Overall, these data are consistent with the fact that the *bc*_1_-complex is indeed needed for survival of bradyzoites, while not every tachyzocidal *bc_1_*-inhibitor also inhibits bradyzoites.

In contrast to tachyzoites, the metabolic phenotype in bradyzoites is not dominated by dihydroorotate dehydrogenase (Fig. 6B, C, and D), as we observed neither the accumulation of carbamoyl-aspartate, dihydroorotate, and aspartate, nor the depletion of pyrimidines. Instead, AMP and NADH accumulate, which in eukaryotes signifies a low cellular energy status (S.-C. Lin and Hardie 2018; X. Fu et al. 2019) that typically induces AMP-activated protein kinase and a glycolytic response. AMPK is a conserved enzyme with a pivotal role in cellular homeostasis: it responds to elevated AMP levels by repressing anabolic activity and by stimulating glucose uptake and glycolysis, β-oxidation, and ketogenesis (Winder and Hardie 1999; S.-C. Lin and Hardie 2018). *T. gondii* tachyzoites express AMPK which serves an important role in metabolic reprogramming during the lytic cycle of tachyzoites (Li et al. 2023). Transcriptomic and proteomic data indicate its presence in bradyzoites (Garfoot et al. 2019) but its function remains unknown. We observed decreased levels of free glucose during all three treatments that are consistent with stimulated glycolysis. We also observe depleted phosphocreatine levels in bradyzoites (Fig. 6) during treatments. In muscle cells, phosphocreatine acts as an ATP buffer system and replenishes ATP pools from ADP via creatine kinase (Kuby, Noda, and Lardy 1954; Perry 1954). It remains to be seen whether bradyzoites possess this mechanism. We also detect elevated levels of acyl-carnitines in BPQ and MMV1028806 treated bradyzoites. These molecules act as shuttles for the mitochondrial import of fatty acids for β-oxidation. However, this pathway has not been shown active and is deemed absent in *T. gondii* (Seeber, Limenitakis et al. 2008, Shunmugam, Arnold et al. 2022). The presence of acyl-carnitines in bradyzoites might reflect import from the host. It is conceivable that their elevation in response to buparvaquone and MMV1028806 indicates compromised functionality of the host *bc*_1_-complex and subsequently accumulating β-oxidation substrates. Indeed, BPQ has a very broad activity across Apicomplexa (Hudson et al. 1985) and kinetoplastids (Croft et al. 1992).

We investigated the functionality of the mETC of tachyzoites and bradyzoites in response to ATQ, BPQ, and MMV1028806 by Mitotracker staining and thin section electron microscopy. All three treatments decrease overall Mitotracker fluorescence, consistent with a specific decrease mitochondrial electron potential (Supplementary Fig. 5). In contrast, pyrimethamine, clindamycin, and DON did not reduce the Mitotracker signal. Pyrimethamine-treated samples displayed an increased Mitotracker-to-DAPI ratio, which can be attributed to diminished DAPI intensity, likely resulting from inhibition of folate-dependent DNA synthesis via DHFR blockade. Clin-treated samples showed no change in mitochondrial morphology or ratio, in line with its delayed-death phenotype. Similarly, DON, which targets glutamine metabolism, did not alter mitochondrial signal or ratio at 24 hours, though morphological changes were observed. Together, these findings confirm that disruption of mitochondrial membrane potential is specific to compounds targeting the electron transport chain, while inhibitors acting on nuclear or metabolic pathways do not induce rapid mitochondrial dysfunction under the conditions tested. ATQ and MMV1028806 but not BPQ also shrink areas of mitochondrial cross-sections indicating a potential contribution to the observed change in overall fluorescence. These differences might reflect distinct mitochondrial morphologies. In agreement with previous observations (Place et al. 2023), we found that bradyzoite mitochondria differ from those in tachyzoites, predominantly occurring as “blobs” rather than tubes or loops, and our observations align with these findings (Supplementary File 6).

The oxygen consumption assays highlight the functional sensitivity of the T. gondii RHΔku80 mETC to pharmacological perturbation (Fig. 5E). As expected, FCCP, an uncoupler of oxidative phosphorylation, markedly increased oxygen consumption, confirming the parasite’s ability to upregulate electron flow when the proton gradient is dissipated. Antimycin A nearly abolished respiration, demonstrating the essential role of complex III in maintaining mitochondrial activity. Likewise, ATQ and BPQ, both *bc1*-complex inhibitors, strongly suppressed oxygen consumption. Notably, MMV1028806 produced a comparable inhibitory effect, suggesting that it also targets complex III or a functionally linked site within the mitochondrial electron transport chain. This observation differs from the findings of (Hayward et al. 2023), who did not report MMV1028806 as an mETC inhibitor in their Seahorse-based screen of the MMV Pathogen Box. A key difference is that our assays were performed at 10 µM, whereas (Hayward et al. 2023) used 1 µM. These results support a mitochondrial mode of action for MMV1028806 under our experimental conditions and underline the value of oxygen consumption assays for detecting dose-dependent effects on parasite respiration.

Our findings strongly suggest that bradyzoite survival is critically dependent on mitochondrial electron transport. We investigated this by comparing the effects of two inhibitors: HDQ, which kills tachyzoites but not bradyzoites, and ATQ, which kills both. Although HDQ enters bradyzoites and reduces their mitochondrial membrane potential, its inability to kill them implies its molecular targets are non-essential in this stage. To pinpoint the vital process, we measured ATP levels and found that only the bradyzocidal compound, ATQ, caused a collapse in bradyzoite ATP. HDQ did not, despite both drugs depleting ATP in tachyzoites. This directly links bradyzoite viability to mitochondrial ATP production. The different outcomes are explained by their distinct targets: ATQ inhibits the *bc1*-complex (McFadden et al. 2000), which is central to ATP synthesis, whereas HDQ blocks other pathways like pyrimidine synthesis and mitochondrial NADH oxidation through DHODH and NDH2. These results also support the notion that the canonical TCA cycle is not the primary source of electrons for bradyzoites (Christiansen et al. 2022). A variety of electron sources supply the electron transport chain (Maclean et al. 2021). The sources that are crucial for ATP production remain to be identified.

Bradyzoites may also utilize glycolysis to generate ATP. This has been suggested to be the case in early, seven-day-old bradyzoites where gliding motility, an ATP dependent process, can be inhibited by withdrawal of exogenous glucose and by the glycolytic inhibitor 2-deoxyglucose. However, in these parasites, also Oligomycin A inhibited motility suggesting a simultaneously essential contribution of the mitochondrial ATP (Fu, Brown et al. 2021). In addition, hexokinase has been shown to be needed for optimal cyst production in chronically infected mice, suggesting an important role of this protein either though its enzymatic activity or other functions (Shukla et al. 2018). It remains to be investigated how mature bradyzoites rely on glycolysis to produce ATP and how this is integrated with mitochondrial ATP production.

Interestingly, despite its profound effect on ATP levels and viability of bradyzoites, ATQ fails to eradicate all *T. gondii* cysts from infected mice and does not reliably prevent toxoplasmosis relapse in AIDS patients and mice (Araujo, Huskinson, and Remington 1991; Kovacs 1992; Ferguson et al. 1994; Katlama, Mouthon, et al. 1996). Underlying reasons for this discrepancy need to be established but may include insufficient bioavailability. Quinolones are known to be poorly soluble in aqueous environments. In addition, the predicted permeability of the blood-brain barriers to either ATQ, BPQ and HDQ is low. Indeed, improved bioavailability of ATQ in nanosuspensions improved efficacies (Shubar et al. 2011). Also, BPQ treatment of mice that were experimentally infected with closely related *Neospora caninum* parasites did target parasites in muscle but not in brain tissue (Müller et al. 2015). A similar conjecture arose when comparing endochin-like quinolones. Here, favorable pharmacokinetics of a variant with lower intrinsic potency likely conferred higher *in vivo* efficacy against cysts (Doggett et al. 2012).

Together, our data illustrate the functional importance of the *bc*_1_-complex for the viability of bradyzoites and encourage the pursuit of *bc*_1_-complex inhibitors as candidates for curative toxoplasmosis treatment. Both, efforts to develop multi-site-directed inhibitors (Doggett et al. 2020; Alday, Nilsen, and Doggett 2022) and re-purposing of respective drugs are promising avenues. However, our data also underline the importance of favorable pharmacokinetics to ensure passage through the blood-brain barrier and the cyst wall.

## Acknowledgements / Funding

We thank Naohiro Hashimoto for sharing the KD3 cells and the Medicine for Malaria Venture for the Pathogen Box and Silvio Bürge for his imaging support.

MB, DM, EP are funded by the Federal Ministry of Education and Research (BMBF) under project number 01KI1715 as part of the "Research Network Zoonotic Infectious Diseases". MB, DM, FS, TH, ML are funded by the Robert Koch Institute.

## Supplements

**Figure S1:**
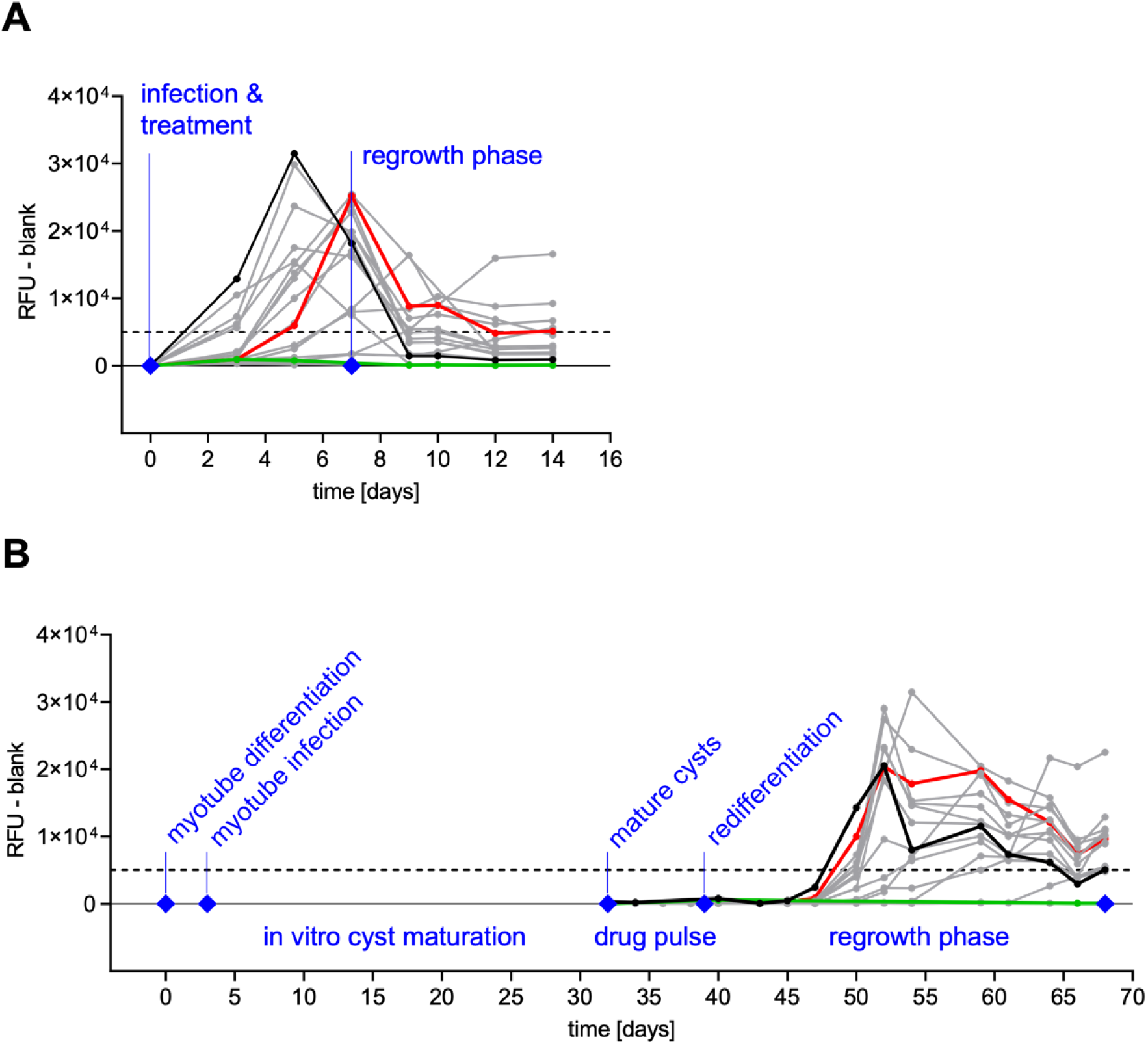
Longitudinal Screening Procedure. All 400 compounds of the MMV Pathogen Box were screened against (A) tachyzoites (n = 4) and (B) in vitro tissue cysts (n = 3). Plotted fluorescence values were background-subtracted and reflect the abundance of parasites per treatment over time (black line = solvent control, green line = inhibitory compound, red line = ineffective compound, dashed line = growth threshold).

**Figure S2:**
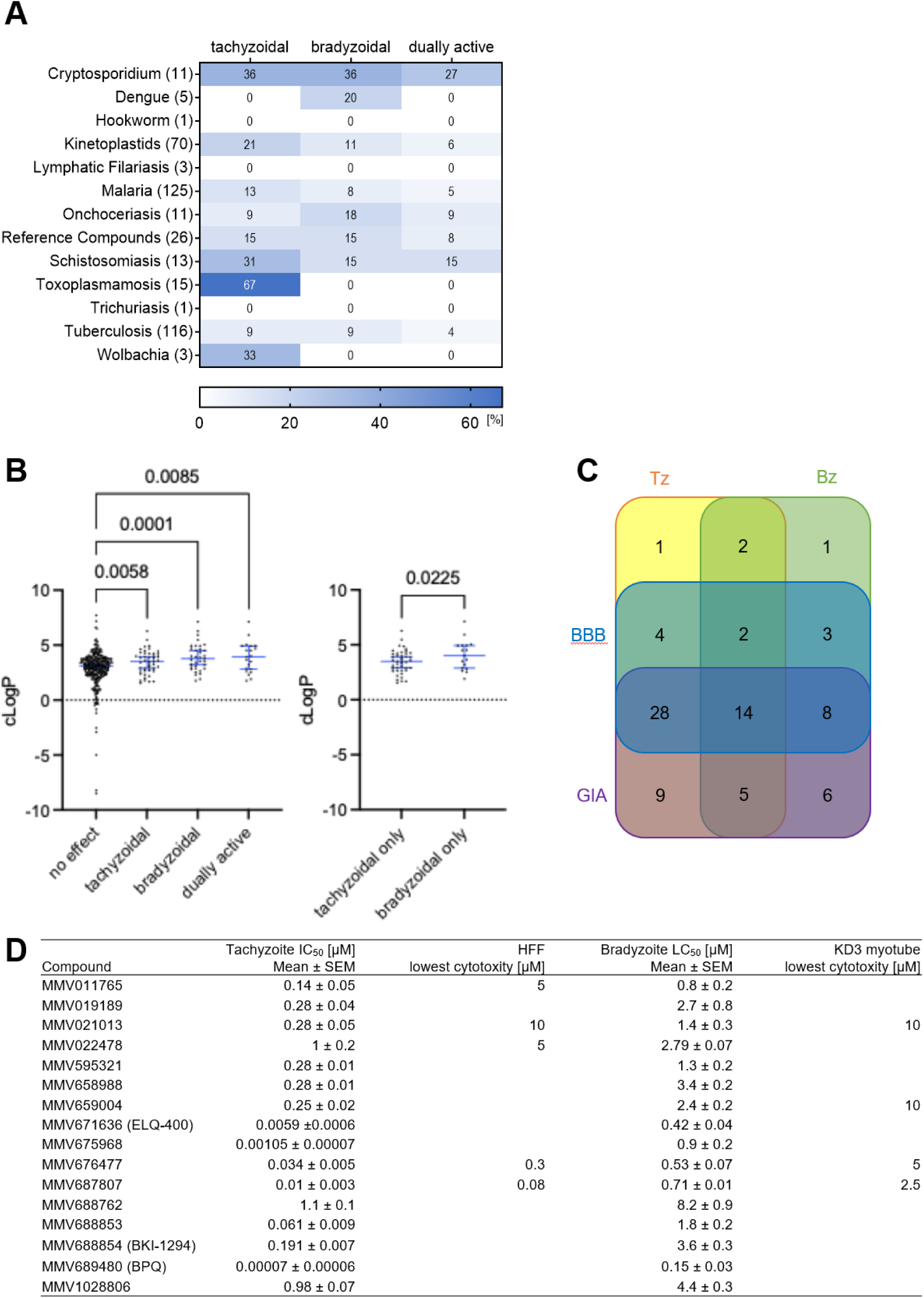
Properties of dually active screening hits. (A) Table showing the relative amount of tachyzocidal, bradyzocidal and dually-active compounds per originally indicated target species. E.g. eleven compounds were indicated by MMV to be active against *Cryptosporidium* and 36% and 27% of these are bradyzocidal and dually active, respectively. In contrast, 67% of the 15 compounds indicated as active against *Toxoplasma* possessed activity against tachyzoites but none were active against bradyzoites. (B) Estimated lipophilicity approximated by predicted partition coefficient suggest that active compounds generally exhibit an increased lipophilicity compared to inactive compounds. (C) A Venn diagram showing intersections of tachyzocidal and bradyzocidal compounds as well as their predicted gastrointestinal absorption capability (GIA) and blood brain barrier (bbb) permeability. (D) Tabulation of compound IC_50_ and LC_50_ data of confirmed active compounds. The calculated IC_50_s and LC_50_s of 16 dually active compounds from figure 2 are summarized in this table. The tachyzoite IC_50_values represent 8 replicates, the LC_50_ values where generated from 6 replicate bradyzoite cultures. Lowest cytotoxicity values were recorded during the primary screen as detailed in the methods section.

**Figure S3:**
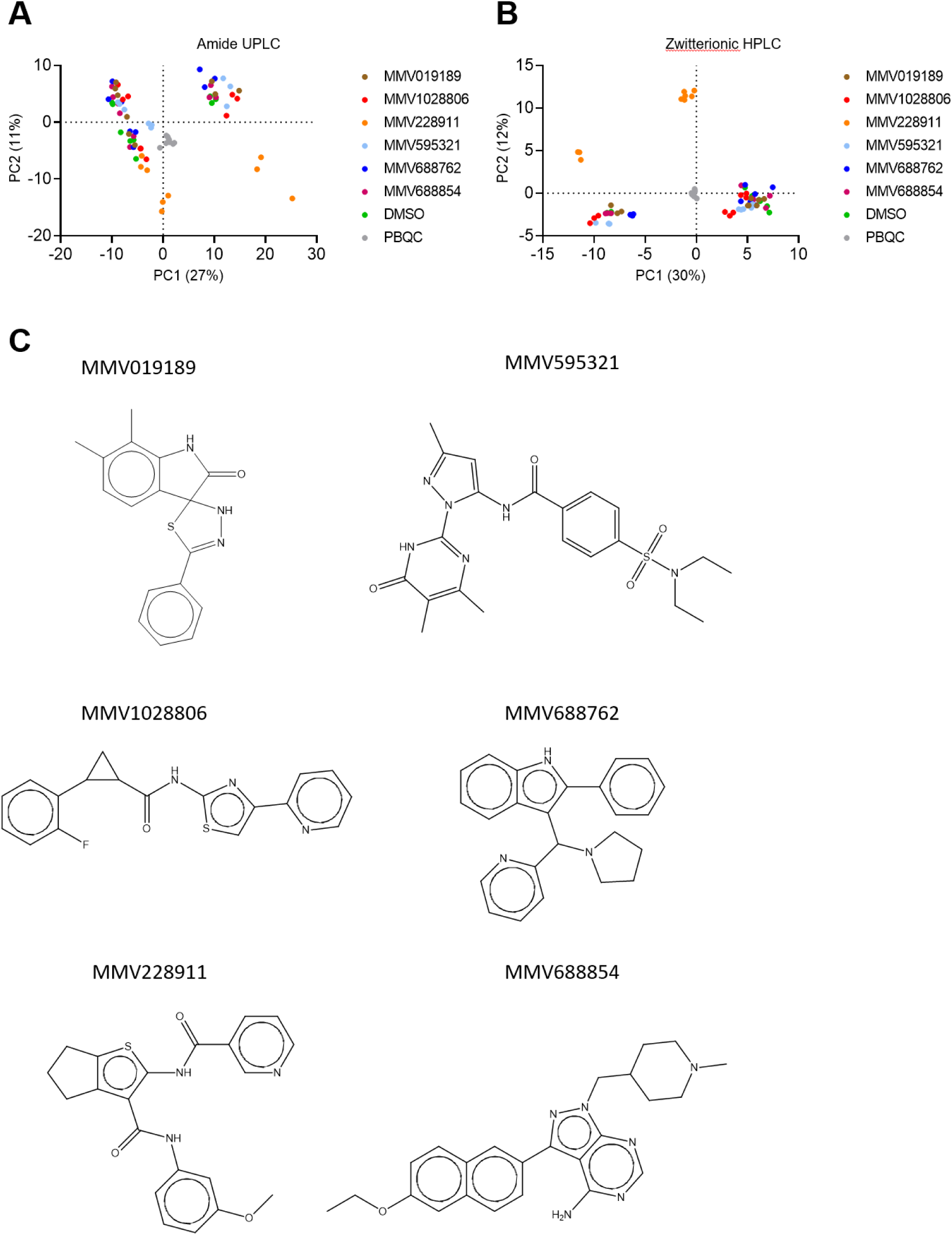
Principal component analysis of LCMS metabolomes of treated parasites using two chromatographic columns. (A) Principal component analysis of metabolites from extracellular tachyzoites that were treated with indicated MMV compounds. Metabolites were separated using BEH-Amide column -HILIC chromatography and detected by mass spectrometry. (B) The same metabolite extracts have been re-analyzed using pHILIC-column HILIC chromatography for separation instead. (C) Chemical structures of dually active and tested compounds as provided by MMV.

**Figure S4:**
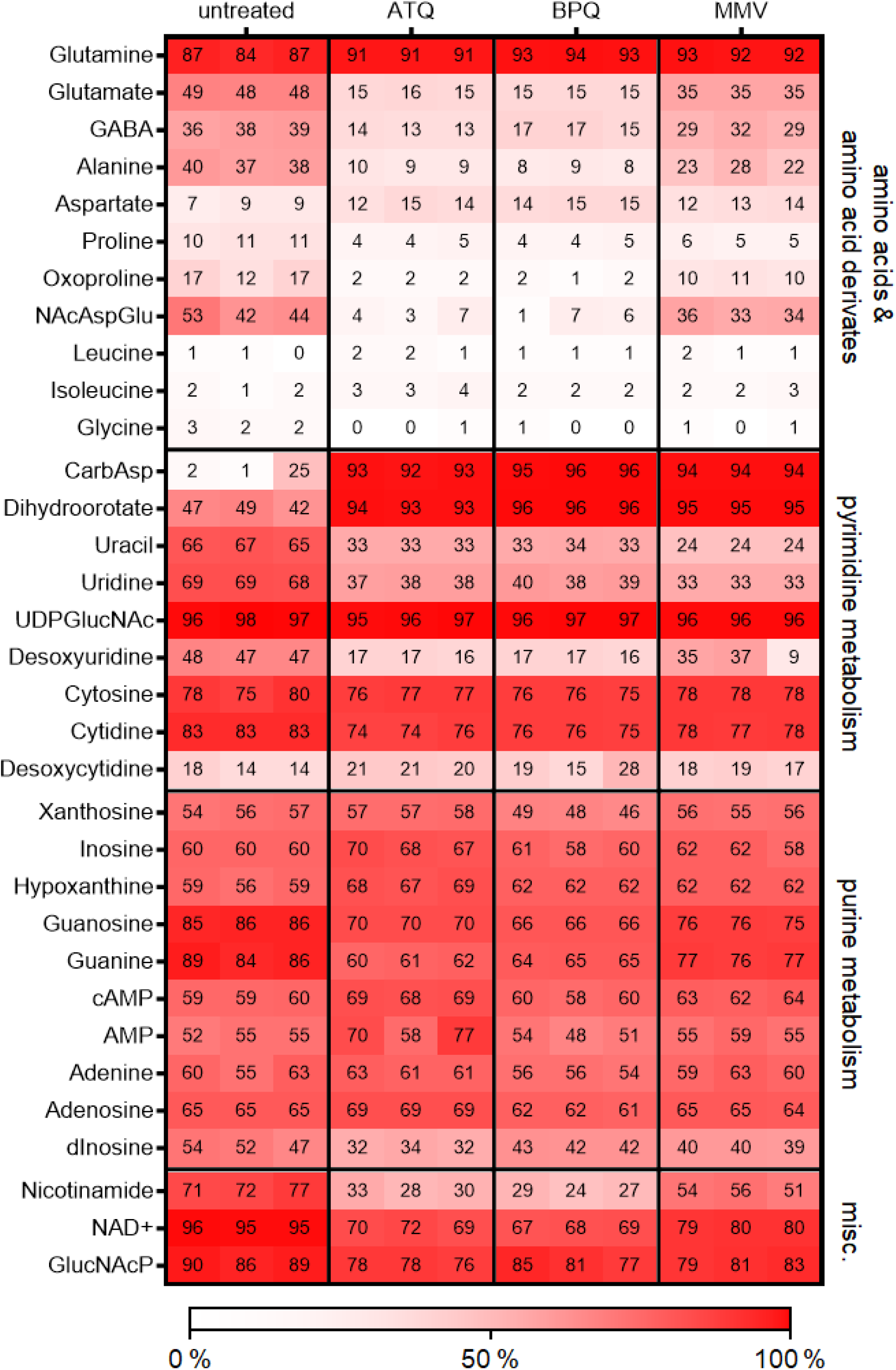
Intracellular tachyzoites exhibit differential ^15^N incorporation after treatment with mETC inhibitors. RHΔku80 were treated with atovaquone, buparvaquone and MMV1028806, labeled with ^15^N-amide-L-glutamine for 3h, followed by their isolation from the HFF cells. The heatmap shows the overall isotopic labeling (100-"M+0") of each replicate.

**Figure S5:**
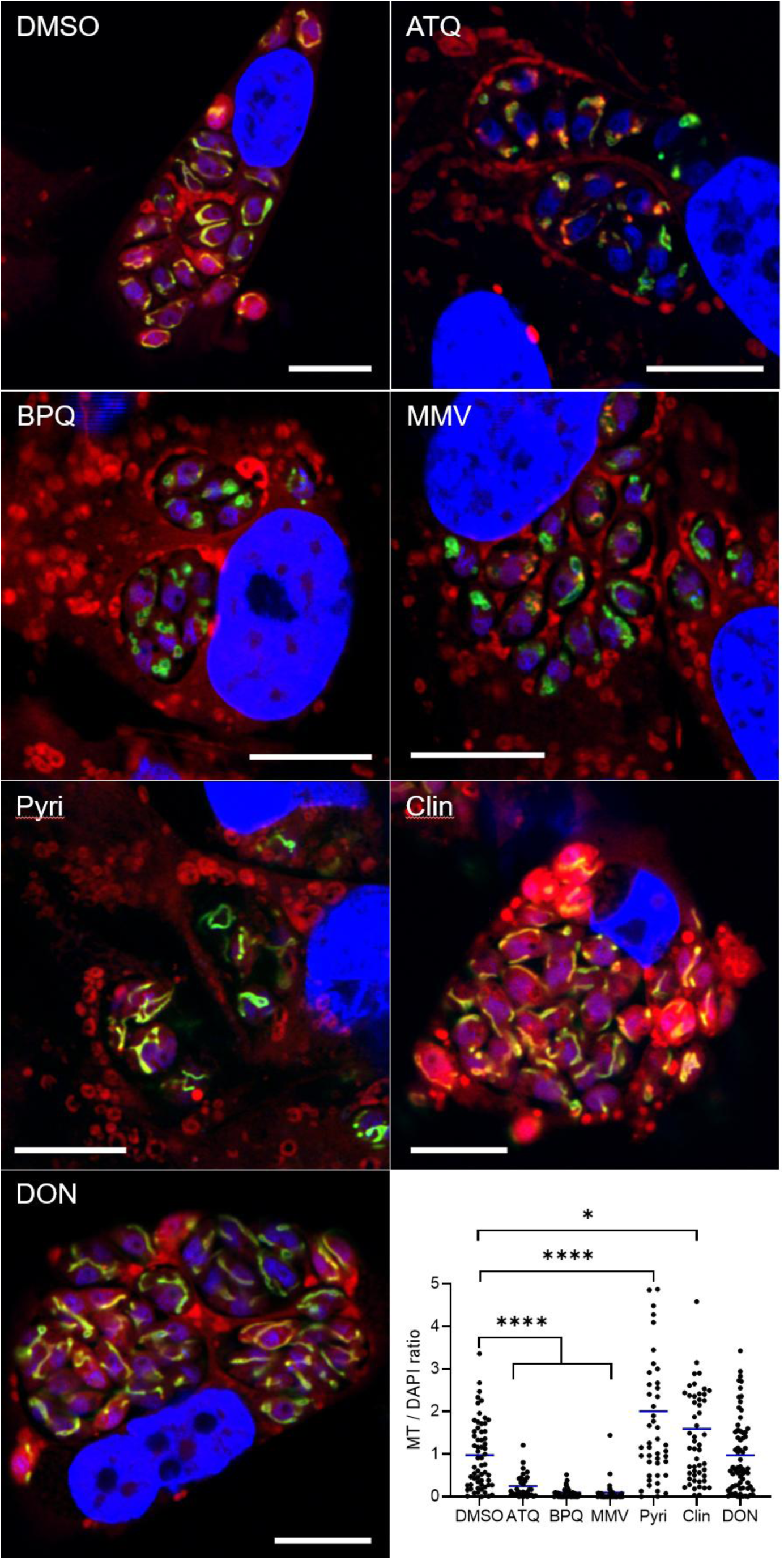
Direct mitochondrial inhibitors reduce Mitotracker signal intensity and MT/DAPI ratio in *T. gondii* tachyzoites. HFF cells were infected with RH-S9 tachyzoites and treated for 24h with DMSO, atovaquone (ATQ), buparvaquone (BPQ), MMV1028806 (MMV), pyrimethamine (Pyri), clindamycin (Clin), and 6-Diazo-5-oxonorleucine (DON). * and **** denote p < 0.05 and p < 0.000 in Mann-Whitney tests).

**Figure S6:**
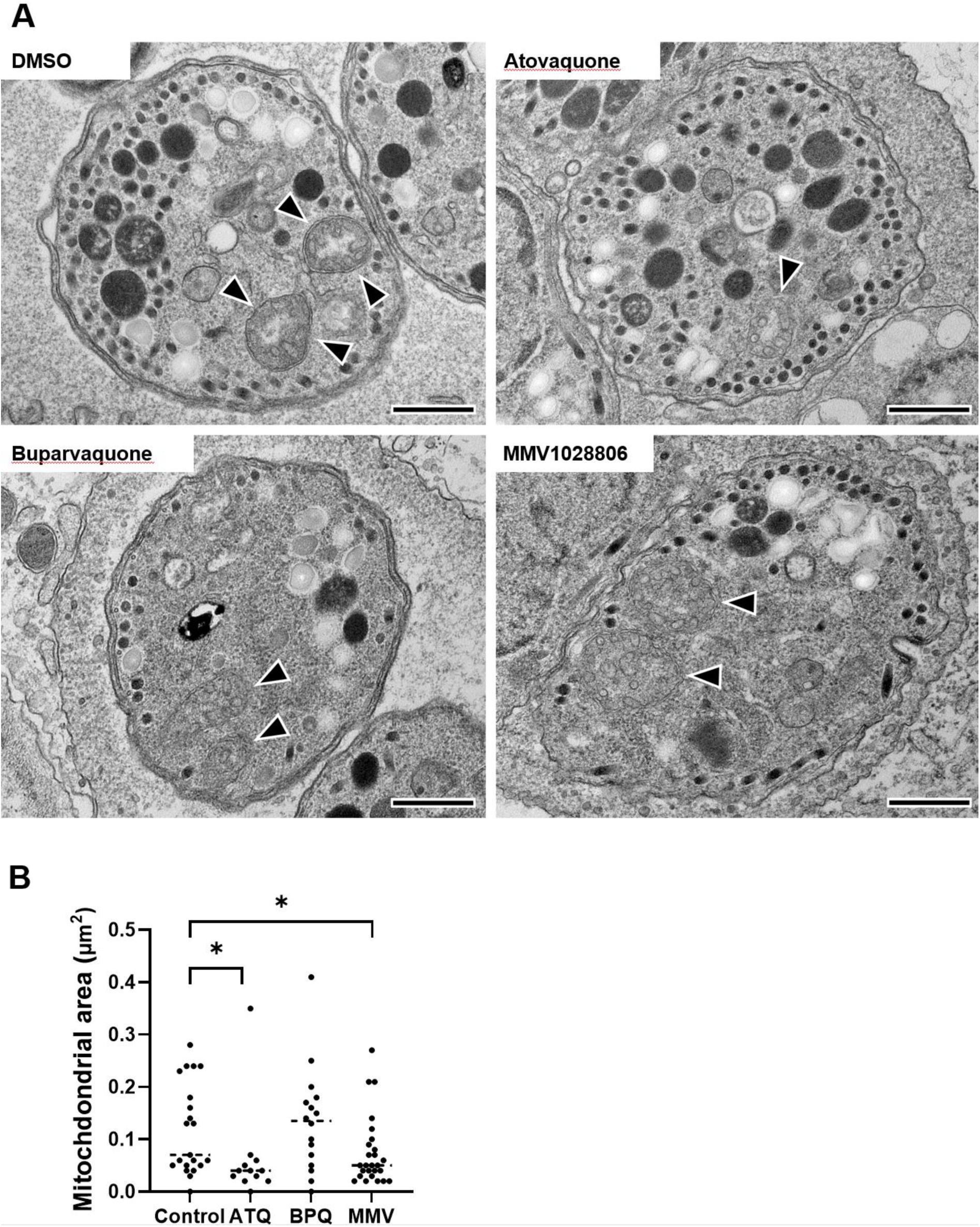
*bc*_1_-complex inhibitors partially reduce mitochondrial profiles of bradyzoites. Thin section electron microscopy of cysts that were treated with atovaquone (ATQ), buparvaquone (BPQ) or MMV1028806 (MMV) for 24 h (A). (B) Measured areas of mitochondrial profiles from 21, 12, 15 and 26 images showing DMSO, ATQ, BPQ and MMV1028806 treated parasites (* denotes p < 0.05 in Mann-Whitney tests). Mitotracker signal appears red, DAPI stain is blue, and the mitochondrial signal originates from GFP. Scale bars represent 10µm. The graph is showing the ratio of the Mitotracker to DAPI signal intensity within each individual parasitophorous vacuole (DMSO n = 61, ATQ n = 32, BPQ n = 35, MMV1028806 n = 36; Pyri n = 58, Clin n = 53, DON n = 68. Blue lines represent the median, two-sided Mann-Whitney test).

**Figure S7:**
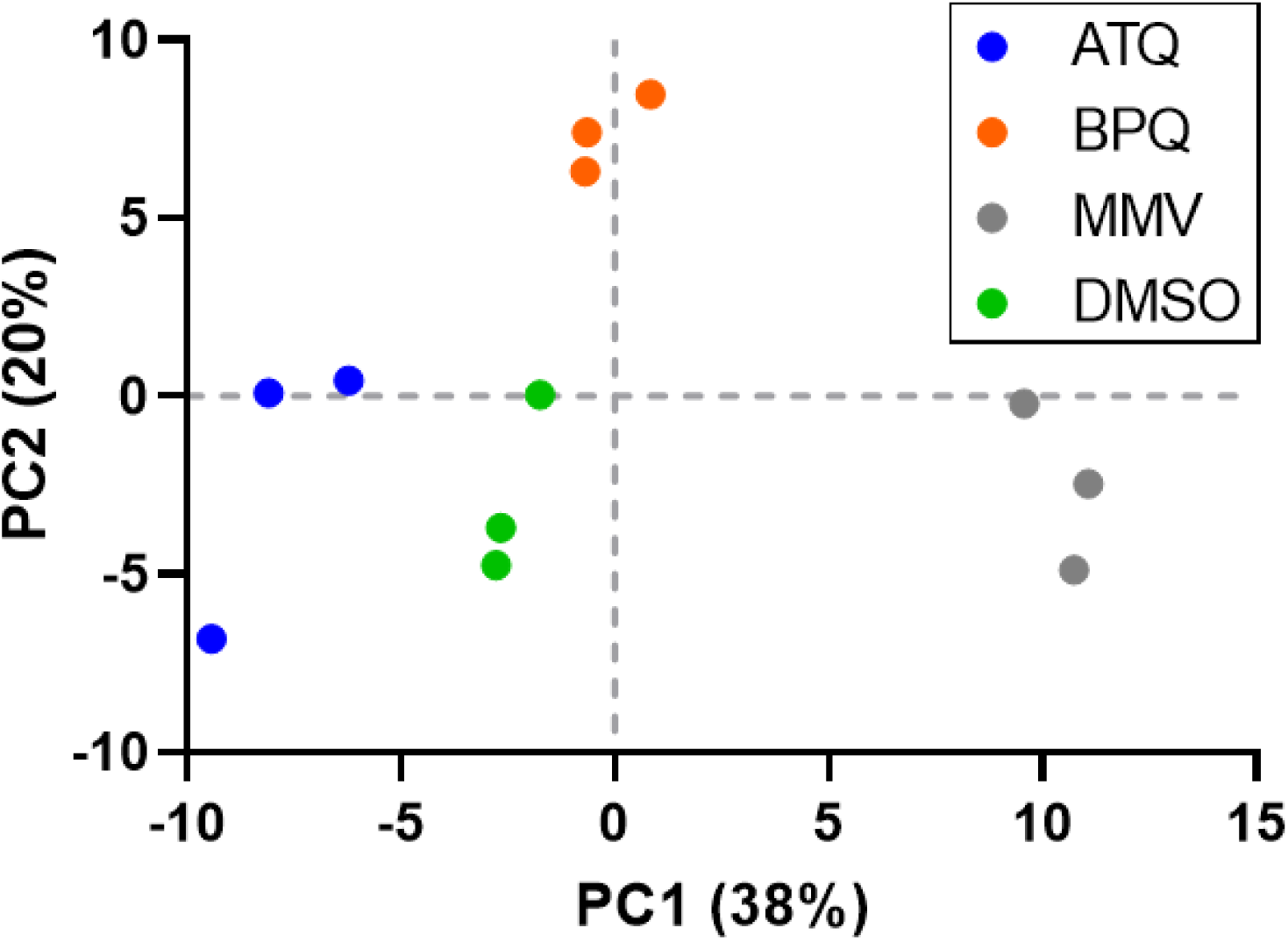
Principal component analysis of bradyzoites treated with atovaquone (ATQ), Buparvaquone (BPQ) and MMV1028806. Four-week-old intracellular cysts were exposed for three hours, purified from their host cells using magnetic beads and analyzed by LCMS.

**Figure S8:**
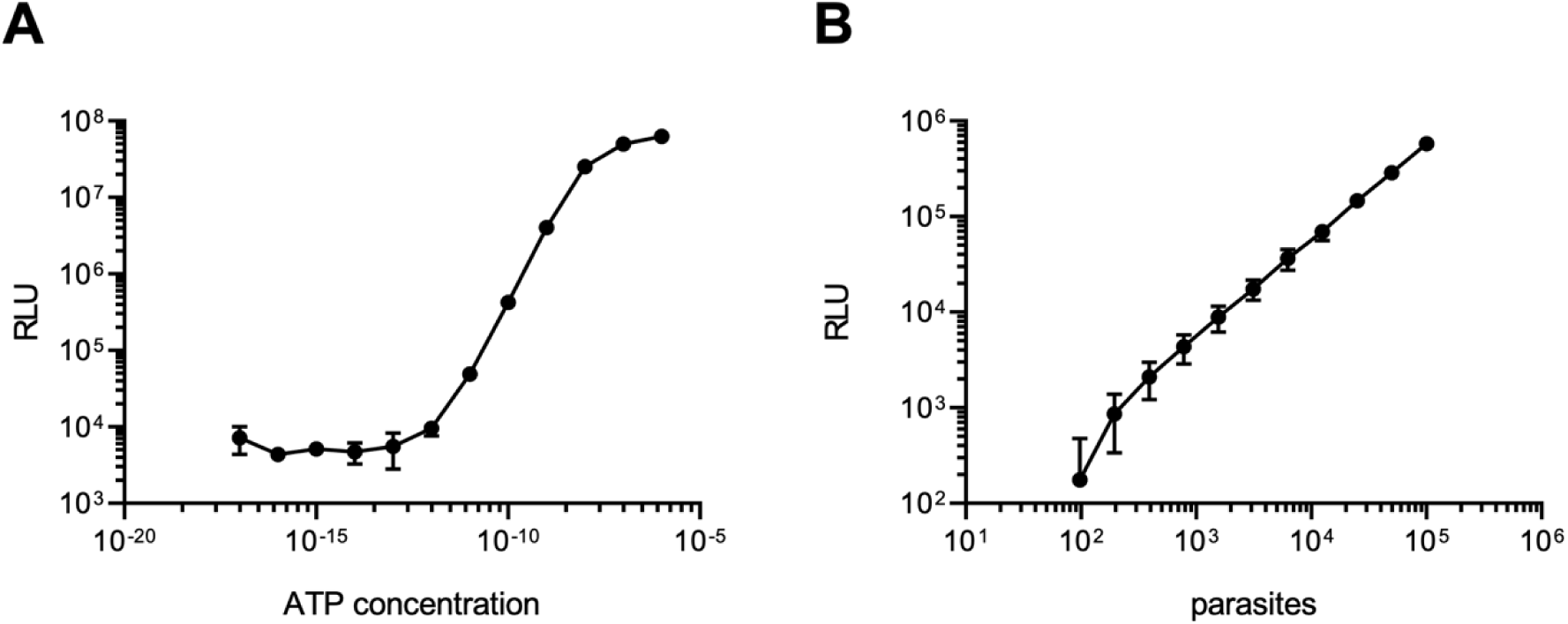
Luminescence-based ATP assay. (A) The linear response of the BacTiter-Glo™ assay has a certain range, depending on the ATP concentration of the sample. The assay is reliable between 100 fM and 1 nM ATP. (B) Increasing numbers of freshly isolated parasites were analyzed to determine a minimal parasite density necessary for steady readouts. 10^5^ parasites per sample were both manageable and accurate.

